# Larger active site in an ancestral hydroxynitrile lyase increases catalytically promiscuous esterase activity

**DOI:** 10.1101/2020.04.06.027797

**Authors:** Bryan J. Jones, Robert L. Evans, Nathan J. Mylrea, Debayan Chaudhury, Christine Luo, Bo Guan, Colin T. Pierce, Wendy R. Gordon, Carrie M. Wilmot, Romas J. Kazlauskas

**Affiliations:** Department of Biochemistry, Molecular Biology and Biophysics and The Biotechnology Institute, University of Minnesota, Saint Paul, Minnesota, United States of America

**Author notes:** Bio-Techne Corporation, Minneapolis, Minnesota, United States of America (Bryan J. Jones). Pace Analytical Services, LLC, Minneapolis, Minnesota, United States of America. Department of Biochemistry, University of Wisconsin, Madison, Wisconsin, United States of America (Debayan Chaudhury). School of Food Science, Shihezi University, Shihezi, Xinjiang, China (Bo Guan).

## Abstract

Hydroxynitrile lyases (HNL’s) belonging to the α/β-hydrolase-fold superfamily evolved from esterases approximately 100 million years ago. Reconstruction of an ancestral hydroxynitrile lyase in the α/β-hydrolase fold superfamily yielded a catalytically active hydroxynitrile lyase, HNL1. Several properties of HNL1 differ from the modern HNL from rubber tree (*Hb*HNL). HNL1 favors larger substrates as compared to *Hb*HNL, is two-fold more catalytically promiscuous for ester hydrolysis (*p*-nitrophenyl acetate) as compared to mandelonitrile cleavage, and resists irreversible heat inactivation to 35 °C higher than for *Hb*HNL. We hypothesized that the x-ray crystal structure of HNL1 may reveal the molecular basis for the differences in these properties. The x-ray crystal structure solved to 1.96-Å resolution shows the expected α/β-hydrolase fold, but a 60% larger active site as compared to *Hb*HNL. This larger active site echoes its evolution from esterases since related esterase SABP2 from tobacco also has a 38% larger active site than *Hb*HNL. The larger active site in HNL1 likely accounts for its ability to accept larger hydroxynitrile substrates. Site-directed mutagenesis of *Hb*HNL to expand the active site increased its promiscuous esterase activity 50-fold, consistent with the larger active site in HNL1 being the primary cause of its promiscuous esterase activity. Urea-induced unfolding of HNL1 indicates that it unfolds less completely than *Hb*HNL (m-value = 0.63 for HNL1 vs 0.93 kcal/ mol·M for *Hb*HNL), which may account for the ability of HNL1 to better resist irreversible inactivation upon heating. The structure of HNL1 shows changes in hydrogen bond networks that may stabilize regions of the folded structure.

## Introduction

Divergent evolution creates superfamilies of enzymes, which share the same protein fold, but differ in substrate specificity or in the type of catalytic activities. The focus of this paper is understanding how evolution creates new catalytic activity during divergent evolution. This question is of interest to evolutionary biologists and also to protein engineers seeking to introduce and optimize new catalytic activities in proteins. Divergent evolution of enzymes to create new catalytic activity is thought to involve intermediate catalytically promiscuous enzymes [1, 2]. Catalytic promiscuity is the ability of enzymes to catalyze additional, chemically distinct reactions besides their primary reaction [3]. Duplication of the genes for these promiscuous enzymes followed by optimization of the promiscuous catalytic activity driven by increased organismal fitness is believed to give rise to enzymes with new primary activities. Support for this notion includes the observation that differing catalytic activities within a superfamily share mechanistic features or transition states and that enzymes within a superfamily often show promiscuous activities that correspond to the primary activities of other enzymes in the super-family [4, 5]. Characterization of resurrected likely ancestral enzymes supports the hypothesis that new functions evolved from ancestors with multiple functions. Reconstructed ancestral enzymes have shown substrate promiscuity [6] and also catalytic promiscuity [7, 8].

Previous work identified three molecular mechanisms that promote catalytic promiscuity in enzymes. First, catalytic promiscuity may have less to do with the enzyme and more with the two reactions being compared. The two reactions may involve similar transition states so the interactions that stabilize the primary reaction also stabilize the promiscuous reaction. In such cases, almost all enzymes catalyzing these reaction types will show promiscuity. For example, the primary function of proteases is amide hydrolysis, but almost all proteases similarly catalyze ester hydrolysis because both reactions involve similar transition states. Second, a catalytically promiscuous enzyme may change its conformation thereby temporarily creating a different enzyme active site structure with different catalytic abilities. For example, a lactonase with promiscuous phosphotriesterase activity catalyzes lactone hydrolysis via a closed conformation, but phosphate triester hydrolysis via an open conformation [9, 10]. The third mechanism for promiscuity is a larger active site with multiple possibilities for interactions between enzyme and transition state. The enzyme active site remains the same, but substrates adopt different orientations within it. For example, phosphonate monoester hydrolase catalyzes promiscuous hydrolysis of sulfate monoesters and other analogs but contains a rigid active site. The varied substrates presumably adopt different orientations within the active site. Catalytic promiscuity correlates with larger active sites and larger polar solvent-accessible surface area [11, 12]. A large active site accommodates a broader range of substrates and allows them to bind in multiple conformations, while a large polar surface allows multiple alternative electrostatic interactions that can stabilize the transition state.

A catalytically promiscuous enzyme may simultaneously use all three of these mechanisms, for example, the lactonase mentioned above also known as paraoxonase I [9]. The enzyme catalyzed hydrolysis of both lactones and phosphate triesters. The first mechanism applies since both hydrolyses have similar negatively charged intermediates and lactonase/phosphotriesterase catalytic promiscuity is common among these enzymes. The lactone hydrolysis intermediates are tetrahedral and the phosphate triester hydrolysis intermediates are pentavalent, but they are nevertheless similar. The second mechanism occurs when different active site conformations enable the two distinct reactions. Finally, the rich catalytic network within the active site enables the third mechanism by promoting multiple reaction pathways by using subsets of active site residues or using them for different roles. In the paraoxonase I case, three amino acid residues (E53, H115, D269) near the active site calcium stabilize the attack of water on the lactone, while only two of these residues (E53, D269) stabilize the attack of water on the phosphotriester.

A reconstructed ancestral hydroxynitrile lyase, HNL1, from the α/β-hydrolase-fold super-family is the focus of this paper [8]. Most enzymes in the α/β-hydrolase-fold superfamily are esterases, which catalyze the hydrolysis of esters, but this superfamily also includes lyases, which catalyze the cleavage of hydroxynitriles and the corresponding reverse addition reaction [13]. HNL1 primarily catalyzes the cleavage of hydroxynitriles, but can also catalyze promiscuous ester hydrolysis. Both reactions involve nucleophilic attack on carbonyl compounds with tetrahedral transition states with partial negative charge on the oxygens. However, the differences in the transition states (acyl enzyme formation versus no acyl enzyme formation, hydrophobic versus polar leaving group) leads to a strong trade-off between esterase and hydroxynitrile lyase catalytic activities. Indeed, it is rare that an esterase catalyzes hydroxynitrile cleavage and vice versa. Thus, the mechanistic basis for the catalytically promiscuous esterase activity of HNL1 remains unclear.

No x-ray crystal structure of a catalytically promiscuous ancestral enzyme has previously been reported, but we expect the mechanistic basis of catalytic promiscuity to be similar to that in modern catalytically promiscuous enzymes. Here we report the x-ray crystal structure of HNL1 and compare the structures and properties of HNL1 and *Hb*HNL (hydroxynitrile lyase from rubber tree, *Hevea brasiliensis*). The larger and more flexible active site of HNL1 as compared to *Hb*HNL allows the ester substrate to adopt the alternative orientation needed for reaction. HNL1, like several other reconstructed ancestral enzymes [14-18], is more thermostable than its modern descendants as measured by melting temperatures. In addition, the denaturation of HNL1 likely only partially unfolds the protein, which allows it to resist the aggregation that causes irreversible inactivation.

## Materials and Methods

### General

Chemicals were bought from commercial suppliers and used without further purification. Racemic mandelonitrile (Sigma-Aldrich, St. Louis, MO) was aliquoted in 10 mL portions and stored at −18 °C. Protein concentrations were determined from the absorbance at 280 nm using calculated extinction coefficients from the ProtParam computational web tool [19]. Protein gels were run using sodium dodecyl sulfate polyacrylamide gradient gels (NuPage 4−12% Bis-Tris gel from Invitrogen) using the BenchMark protein ladder (Invitrogen, 5 μL/lane) as a standard. Nickel affinity chromatography resin (NiNTA, Qiagen) was used to purify HNL1 containing a 6-His tag expressed in *Escherichia coli*. This chromatography resin was regenerated according to Qiagen protocol. ^1^H NMR spectra were run at 400 MHz in deuterochloroform.

2-Hydroxy-2-(6-methoxynaphthalen-2-yl) acetonitrile, **1**. Synthesis of **1** was adapted from Bhunya et al. [20]. NaCN (∼3.9 g) was added to a solution of 6-methoxy-2-naphthaldehyde (1.0 g) in acetic acid (15 mL). CAUTION! This reaction generates hydrogen cyanide which is toxic, see handling guidelines [21]. The reaction was carried in the closed fume hood at room temperature (22 °C) for two weeks. The reaction mixture was neutralized using NaHCO_3_ until ∼pH 5 and extracted twice with ether/dichloromethane. The organic extract was dried using Na_2_SO_4_ and dried by rotary evaporation. The purity was about 97% based on HPLC. The ^1^H-NMR in CDCl_3_ matched the previously reported spectrum [22] and showed no remaining starting material.

### Cloning, gene expression, and protein purification

#### Ancestral enzyme reconstruction

Ancestral enzyme reconstruction of HNL1 [8, 23] started with a Maximum Likelihood phylogenetic tree built using the software tool RAxML [24]. The tree contained 1,285 α/β hydrolase-fold enzyme sequences between 30% and 99% identical and was used to infer ancestral sequences using the same software. This clade included plant hydroxynitrile lyases (HNLs) from *Hevea brasiliensis* and *Manihot esculenta*, as well as plant esterases from *Nicotiana tabacum* and *Rauvolfia serpentine* (UniProtKB: P52704, P52705, Q6RYA0, Q9SE93 respectively). HNL1 is the reconstruction at the node that is the last common ancestor of the HNLs from *Hevea brasiliensis* (*Hb*HNL), *Manihot esculenta* (*Me*HNL), and *Baliospermum montanum* (*Bm*HNL). The gene for the ancestral enzyme HNL1 was synthesized by GenScript (Piscataway, NJ) and subcloned into a pET21a(+) vector at *Nde*I and *Xho*I restriction sites resulting in an upstream T7 promoter and lac operator and a C-terminal six His-tag. Fidelity of cloning and gene synthesis was confirmed by DNA sequencing of the gene (ACGT, Wheeling, IL). Genes for the modern HNLs, *Hb*HNL and *Me*HNL were cloned and expressed in the same way.

#### Site-directed mutagenesis

The HNL1-esterase variant was created by three single-amino-acid substitutions in the gene for HNL1 added stepwise (Thr11Gly, Lys236Gly, Glu79His) in the order listed using the QuickChange II method (Agilent Technologies) according to the manufacturer’s instructions. The residues at three positions in HNL and *Hb*HNL (121,178 and 146) were interchanged by site-directed mutagenesis stepwise in the order listed using the Q5 site-directed mutagenesis method (New England BioLabs, Inc.) according the manufacturer’s instructions. S1 Table lists the primer pairs used for the polymerase chain reaction step of this site-directed mutagenesis. A typical procedure for the Q5 method is as follows. The polymerase chain reaction (total volume 25 µL) used typically 15-20 ng template DNA, 0.5 µM each of forward and reverse primers, 12.5 µL of 2X Q5 Hot Start High-Fidelity Master Mix and nuclease free water. The temperature program was: initial denaturation at 98 °C for 2 minutes, followed by 25 cycles of denaturation at 98 °C for 10 seconds, annealing at a temperature between the melting temperatures of the forward and reverse primers for 20 seconds, and extension at 72 °C for 3.5 minutes. The final step concluded with a 10-min extension at 72 °C followed by cooling to 4 °C for storage. 1 µL-2.5 µL of PCR reaction was visualized on a 1% agarose gel to confirm the presence of linear PCR product. For the Q5 site-directed mutagenesis method the PCR product was then treated with the kinase-ligase-*Dpn*I enzyme mix in a 10 µL reaction containing 1 µL of KLD mix, 5 µL of KLD Buffer, 1-2 µL of PCR product and 2-3 µL of nuclease free water to remove the template DNA as well as to re-circularize the linear PCR product for more efficient transformation. *E. coli* DH5α cells were transformed with the mutated plasmids and allowed to grow overnight at 37 °C. The plasmid DNA was isolated via miniprep (Qiagen) and sequenced prior to initiating the next mutation.

#### Protein expression and purification

HNL1 and its variants expressed well in *E. coli* BL21 and purification by Ni-affinity chromatography yielded high purity protein. Lysogeny broth media containing ampicillin (100 μg/ mL, LB-amp, 5 mL) was inoculated with a single bacterial colony from an agar plate and incubated in an orbital shaker at 37 °C and 200 rpm for 15 h to create a pre-culture. A 1-L baffled flask containing terrific broth (TB)-amp media (250 mL) was inoculated with this pre-culture. This culture was incubated at 37 °C and 250 rpm for 3-4 h until the absorbance at 600 nm reached 1.0 and transferred to 17 °C and 200 rpm for 1 h to cool. Isopropyl β-D-1-thiogalactopyranoside (IPTG, 0.75-1.0 mM final concentration) was added to induce the protein expression, and cultivation was continued for 20 h. The cells were harvested by centrifugation (8000 rpm, 10 min at 4 °C), resuspended in buffer A (10-20 mM imidazole, 50 mM NaH_2_PO_4_, 300 mM NaCl, pH 7.2, 20 mL) and disrupted by sonication (400 W, 40% amplitude for 5 min). The tube was centrifuged (4 °C, 12 000 rpm 45 min) and the supernatant was loaded onto a column containing NiNTA resin (1 mL, Qiagen) pre-equilibrated with buffer A (10 mL). The column was washed with buffer A (50 mL) followed by buffer B (50-60 mM imidazole, 50 mM NaH_2_PO_4_, 300 mM NaCl, pH 7.2, 50 mL). The His-tagged protein was eluted with elution buffer (250 mM imidazole, 50 mM NaH_2_PO_4_, 300 mM NaCl, pH 7.2, 15 mL) and concentrated to ∼1 mL with an Amicon ultrafiltration centrifuge tube (10 kDa cutoff). The imidazole-containing elution buffer was exchanged by four successive additions of BES buffer (5 mM *N,N*-bis(2-hydroxyethyl)-2-aminoethanesulfonic acid, pH 7.0, 10 mL) followed by concentration to 1 mL by ultrafiltration. A 250-mL culture typically yielded 15-35 mg of protein. SDS-PAGE indicated a molecular weight of ∼30 kDa in agreement with the predicted weight of 31.1 kDa. S1 Fig shows the SDS PAGE gel of HNL1 and HNL1 esterase catalysis variant.

### Enzyme assays

Enzyme activity was monitored at ambient temperature in triplicate for 10 min using a microplate reader. The rates were corrected for any spontaneous reaction.

One hydroxynitrile lyase assay, Fig 1, monitored the formation of 6-methoxy-2-naphthaldehyde (ε_315 nm_ = 25,000 cm^-1^ M^-1^) from cyanohydrin **1** at 315 nm. The reaction mixture (100 μL, path length 0.29 cm) in a 96-well UV microtiter plate contained 2-10 μg enzyme, 50 mM sodium citrate-phosphate buffer (pH 5.0), 450 μM of 1 and 5 vol% acetonitrile to dissolve the substrate.

**Fig 1.**
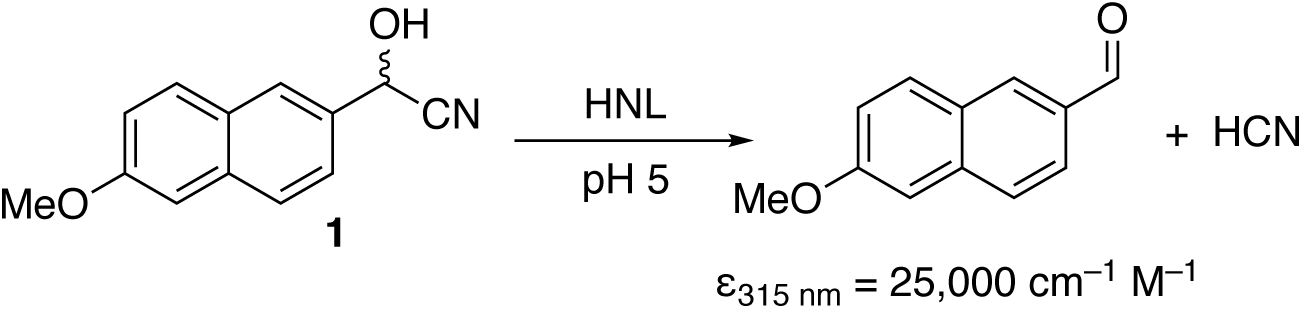
Hydroxynitrile lyase-catalyzed cleavage of cyanohydrin 1 yields an aldehyde that absorbs light strongly at 315 nm.

Another HNL assay monitored the formation of benzaldehyde (ε_280 nm_ = 1380 cm^-1^M^-1^) from racemic mandelonitrile at 280 nm. The reaction mixture (200 μL; path length 0.58 cm) in a 96-well UV microtiter plate contained 1-10 μg enzyme, sodium citrate buffer (50-54 mM, pH 5.0), 0-32 mM mandelonitrile. The slope of increase in absorbance versus time was measured in triplicate, fit to a line using linear regression, and corrected for spontaneous cleavage of substrate with blank reactions lacking protein.

Ester hydrolysis activity was measured at 404 nm using *p*-nitrophenyl acetate (*p*NPAc), which releases the yellow *p*-nitrophenoxide. The reaction mixture (100 μL; path length 0.29 cm) contained 0.03-2.4 mM *p*NPAc, 7-8% vol % acetonitrile, 5 mM BES buffer, pH 7.0, and up to 15 μg enzyme. The slope of increase in absorbance versus time was measured in triplicate, fit to a line using linear regression, and corrected for spontaneous hydrolysis of *p*NPAc with blank reactions lacking protein. The extinction coefficient used for calculations (ε_404 nm_ = 11,400 cm^-1^M^-1^) accounts for the incomplete ionization of *p*-nitrophenol at pH 7.0. For some esterase assays, the pH was 7.2 and an extinction coefficient of 16,600 cm^-1^M^-1^ was used. For steady-state kinetic measurements, the enzyme concentration was determined by average absorbance at 280 nm measured in duplicate and normalized by subtracting a buffer blank. The enzyme concentrations of purified HNL1 and the HNL1 variants in the assay solution were both ∼1.75 mg/mL (56 µM). The k_cat_ and K_M_ were determined using a non-linear fit of the experimental data to the Michaelis-Menten equation using the solver program in Microsoft Excel or using the statistical program R [25, 26]. (Huitema & Horsman, 2019). S2 Fig shows typical experimental data and fit.

#### Measuring catalytic promiscuity

To quantitatively compare the catalytic promiscuity of different enzymes, we define below a measure of promiscuity based on comparison of two substrates in analogy to substrate specificity. This focus on specific substrates facilitates identification of specific molecular features that favor one reaction over the other but suffers the disadvantage that some of these differences will be substrate-specific instead of catalytic-reaction specific. Other researchers have defined catalytic promiscuity indices based on the number of reactions catalyzed [27], but these measures do not account for changes in the relative rates of these reactions. Other promiscuity measures only apply to substrate promiscuity [28] or are based only on structure, not catalytic properties [29].

Selectivity of an enzyme is the ratio of its catalytic efficiency for a fast reaction over a slow reaction, eq 1. In most cases, this measure is used to compare two substrates. For example, enantioselectivity measures the ability of an enzyme to discriminate between the two enantiomers. A selectivity of 1 corresponds to a non-selective reaction, while large values correspond to high selectivity.

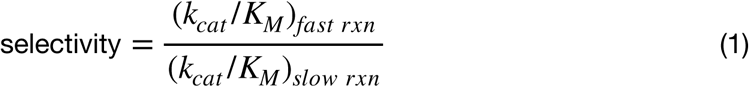

Promiscuity is the opposite of selectivity; it is the ability to efficiently catalyze multiple reactions. We define promiscuity for any pair of comparison reactions, P, as the inverse of selectivity, eq. 2.

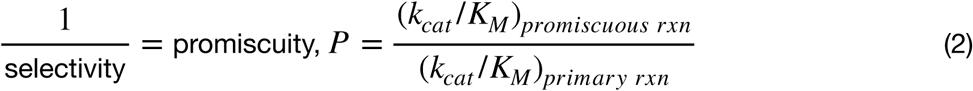

A promiscuity of 1 corresponds to equal catalytic efficiency of both reactions. Values less than one correspond to lower promiscuity, while values more than one correspond to a switch in the primary reaction. As for selectivity, promiscuity is defined by specifying a pair of reactions. Substrate promiscuity is defined by a pair of substrates, while reaction promiscuity is defined by a pair of reactions where the substrate must be specified as well.

### Thermostability

Stability was assessed by three methods. First, irreversible inactivation was evaluated by heating enzyme aliquots in a water bath at a set temperature (70-95 °C) for 15 or 30 min and then cooling on ice for at least 15 minutes. The heated and unheated enzyme samples were assayed for HNL activity using mandelonitrile as described above. Stability is expressed as the percent activity retained after heating.

Second, the melting point (T_m_) of the enzyme also indicated stability. Microplates (384-well) containing 12 replicates of each sample (10 µL containing 300 ng protein, 50 mM sodium PIPES (piperazine-N,N′-bis(2-ethane sulfonic acid), 100 mM NaCl) were heated at 1 °C /min while monitoring the intensity and the lifetime of the intrinsic fluorescence using a NovaFluor PR instrument (Fluorescence Innovations, Inc., Minneapolis, MN). Melting curves were fit to the lifetime and intensity data and yielded similar T_m_ values; an average value is reported.

We used the following approximation to convert differences in T_m_ to differences in Gibbs energies of unfolding. A comparison of pairs of homologous proteins showed that the melting temperatures, T_m_, were 31.5 °C higher and the Gibbs energies of unfolding 8.7 kcal/mol higher for the proteins from thermophiles as compared to the protein from mesophiles [30]. From this comparison, we can derive a rule of thumb that stabilizing a protein by 1 kcal/mol will increase the melting temperature by ∼3.6 °C.

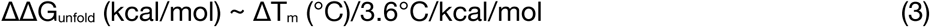

The third measure of stability was urea-induced unfolding. Proteins *Hb*HNL and HNL1 were placed in increasing concentrations of urea (0-7.2 M), and the resulting decrease in tryptophan fluorescence was measured using a SpectraMax GEMINI XS plus-384 plate reader with an excitation wavelength of 278 nm and an emission wavelength of 329 nm. Enzyme (20 μL) with a concentration of 0.1-0.3 mg/mL was placed in each well of a 96-well microtiter plate, along with 180 μL of different mixtures of buffer (5 mM BES buffer pH 7.2) and 8 M urea solution for a total of 200 μL. The plate was equilibrated at 4 °C for 24 hours, then warmed to room temperature before measuring the fluorescence. The relative concentrations of the folded or native conformation, *N*, and unfolded or denatured state, *U*, were determined by comparing the fluorescence at low urea concentrations (completely folded) and high urea concentrations (completely unfolded). The equilibrium ratio of folded and unfolded forms at a particular denaturant concentration yields the free energy of protein unfolding, Δ*G*_*unfold*_, at that denaturant concentration, eq 4,

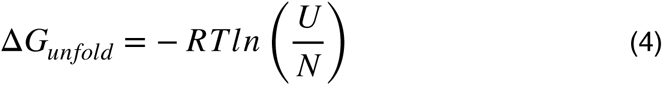

where R is the gas constant and T is the temperature. A plot of this free energy versus urea concentration yields a straight line of the form in eq 5,

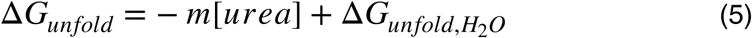

where *m* is the slope and represents the sensitivity to denaturation by urea. Δ*G*_*unfold,H2O*_ is the y-axis intercept and represents the free energy of unfolding in the absence of denaturant [31]. The linear model in the statistical software R [25] was used to fit the variation of Δ*G*_*unfold*_ with urea concentration to a line.

### Crystallization and x-ray data collection

The High Throughput Crystallization Center at the Hauptman-Woodward Institute (Buffalo, NY, USA) screened crystallization conditions using droplets under oil. Droplets of protein solution (200 nL, 10 mg/mL in 5 mM BES, pH 7.2) were mixed with an equal volume of crystallization solution and equilibrated at 24 °C under oil. Crystals formed after three weeks in several conditions including ammonium citrate (2.0 M) in bis-tris propane (0.1 M, pH 7). Adapting similar conditions to a 24-well hanging drop vapor diffusion resulted in crystals suitable for x-ray data collection. Protein solution (1 µL, 7 mg protein/mL in 5 mM BES, pH 7.2) was combined with reservoir solution (1 µL, 1.1 M ammonium citrate, 50 mM bis-tris propane, pH 6.8) and equilibrated at 19 °C. Crystals were harvested at day 20.

Before data collection, crystals were transferred to a cryosolution consisting of a well solution with 5% glycerol for 5-10 s and flash-cooled in liquid nitrogen. The x-ray diffraction data were collected at 100 K on beamline 23 ID-B (Eiger 16M detector) at Advanced Photon Source at Argonne National Labs, Lemont, IL.

### Structure determination

The data sets were integrated with *XDS* [32] and scaled using Aimless from the *CCP*4 suite [33, 34]. The structure was determined by molecular replacement with *Phaser MR* [35] using the coordinates of *Hb*HNL with pdb id 1yb6 [36], having 81% sequence identity to HNL1. Search models were derived from these coordinates by removing side chains and loop regions that differ using structure-based sequence alignment. Molecular replacement using this truncated model gave one solution that was clearly above the background, with scores: RFZ=3.3 PAK=0 LLG=5261 TFZ==29.2. Four molecules were found in the asymmetric unit. A full structural model could be built into the electron density obtained from these phases.

Phase improvement was made with *REFMAC5* [37]. Model building was done with *Coot* [38]. Residues 2-260 were modeled in all HNL1 chains, with some also including a portion of the C-terminal linker to the 6xHis tag. The 6xHis tag and methionine 1 were unable to be modeled in any of the chains. The final refinement run with *REFMAC*5 included TLS refinement [39]. *MolProbity* [40] was used to analyze sterics and geometry during the refinement process.

The final model contains four polypeptide chains of HNL1, six glycerol molecules (including one in the active site of each chain), and 192 water molecules. The crystallographic data for HNL1 have been deposited in the Protein Data Bank (pdb id 5tdx). Figures of the structure were prepared with *PyMOL* (https://www.pymol.org).

#### Normalization of B-factors

The average and standard deviation of the B-factors for the Cα atoms were used for the normalization of B-factors. For each Cα B-factor, the average was subtracted and the difference was divided by the standard deviation [41]. This Z-score value identifies regions that differ from the average value in units of standard deviation. For example, the average B-factor for the Cα atoms of chain A in 5tdx is 32.4 Å^2^ with a standard deviation of 7.4 Å^2^. The Cα atom for Ala2 has a B-factor of 61 Å^2^, which corresponds to Z-score of (61-32.4)/7.4 = 3.9. This Cα carbon is 3.9 standard deviations more flexible than an average residue in this chain. High flexibility at the N- and C-terminal ends of a protein is common.

#### Volume of active site

A tunnel leading from the active site to the enzyme surface was defined for each enzyme using the web tool, MOLEonline 2.0 (https://mole.upol.cz; [42], with default settings. The same tunnel was compared for each enzyme. An innermost part of this tunnel corresponds to the active site. The start of the active site was defined relative to the start of the tunnel in *Hb*HNL (pdb id 1yb6) and the end of the active site was defined as 10.5 Å from the start because *Hb*-HNL shows a pinch point in the tunnel near this point. See text for details of this choice as the end of the active site. MOLEonline calculates tunnel volumes by summing the multiple, constituent frustums (parallel truncations of right cones). At a given cross-section the three closest atoms in the tunnel define the radius r of a circle forming one end of each frustum. MOLEonline set the number and height of each frustum depending on the tunnel geometry. The tunnel volume is the sum of the volumes of the constituent right conical frustums, eq 6, where r_1_ and r_2_ are the radii of the circles forming the ends of each frustum and h is the height of each frustum.

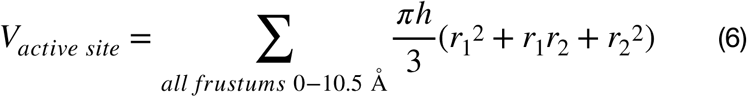

#### Docking of *p*-nitrophenyl acetate

SwissDock [43] was used to dock *p*NPAc to the active sites of *Hb*HNL (pdb id 1yb6), HNL1 (pdb id 5tdx), and SABP2 (pdb id 1y7i). The region of interest was centered at the Oγ of the active site serine and extended 16 Å along the x, y, and z axes. The protein side chains within 5 Å of *p*NPAc were set to flexible during docking.

## Results

### Comparison of properties of HNL1 and modern HNLs

To test the hypothesis that ancestral enzymes are more catalytically promiscuous that their modern descendants, we previously resurrected a putative ancestral hydroxynitrile lyase, HNL1, within the α/β-hydrolase fold superfamily [8]. This ancestral enzyme separates a small group of hydroxynitrile lyases from the vast number of esterases in this superfamily. We compared the substrate specificity and catalytic promiscuity of this ancestral HNL1 to three of its modern descendants whose x-ray crystal structures have been solved: HNLs from rubber tree (*Hevea brasiliensis, Hb*HNL), cassava (*Manihot esculenta, Me*HNL), and wild castor (*Baliospermum montanum, Bm*HNL). HNL1 and *Hb*HNL share 81% identical amino acid residues over a comparison of 257 residues. This comparison excludes the His tag and linker. HNL1 shares 77% identical amino acids residues with *Me*HNL and 67%with *Bm*HNL.

#### Substrate specificity and catalytic promiscuity of HNL1

HNL1 favors larger substrates as compared to *Hb*HNL, Fig 2. This generalization applies to both lyase-type reactions and ester hydrolysis. The lyase type reactions tested are the cleavage of various cyanohydrins and the cleavage of 2-nitro-1-phenylethanol. This second cleavage is a promiscuous reaction and is the reverse of a nitro-aldol addition. Substrate molecular weight approximates substrate size on the x-axis, while the natural logarithm of the ratio of HNL1-catalyzed rate over the *Hb*HNL-catalyzed rate compares the two rates. The logarithm avoids squeezing ratios less than one into the region between 0 and 1, but instead extends these ratios to negative values. A linear fit of the data yields a moderate r^2^ value of 0.61, indicating that differences in the molecular weight of the substrate account for 61% of the relative rate variation. The positive slope indicates that HNL1 favors larger substrates as compared to *Hb*HNL.

**Fig 2.**
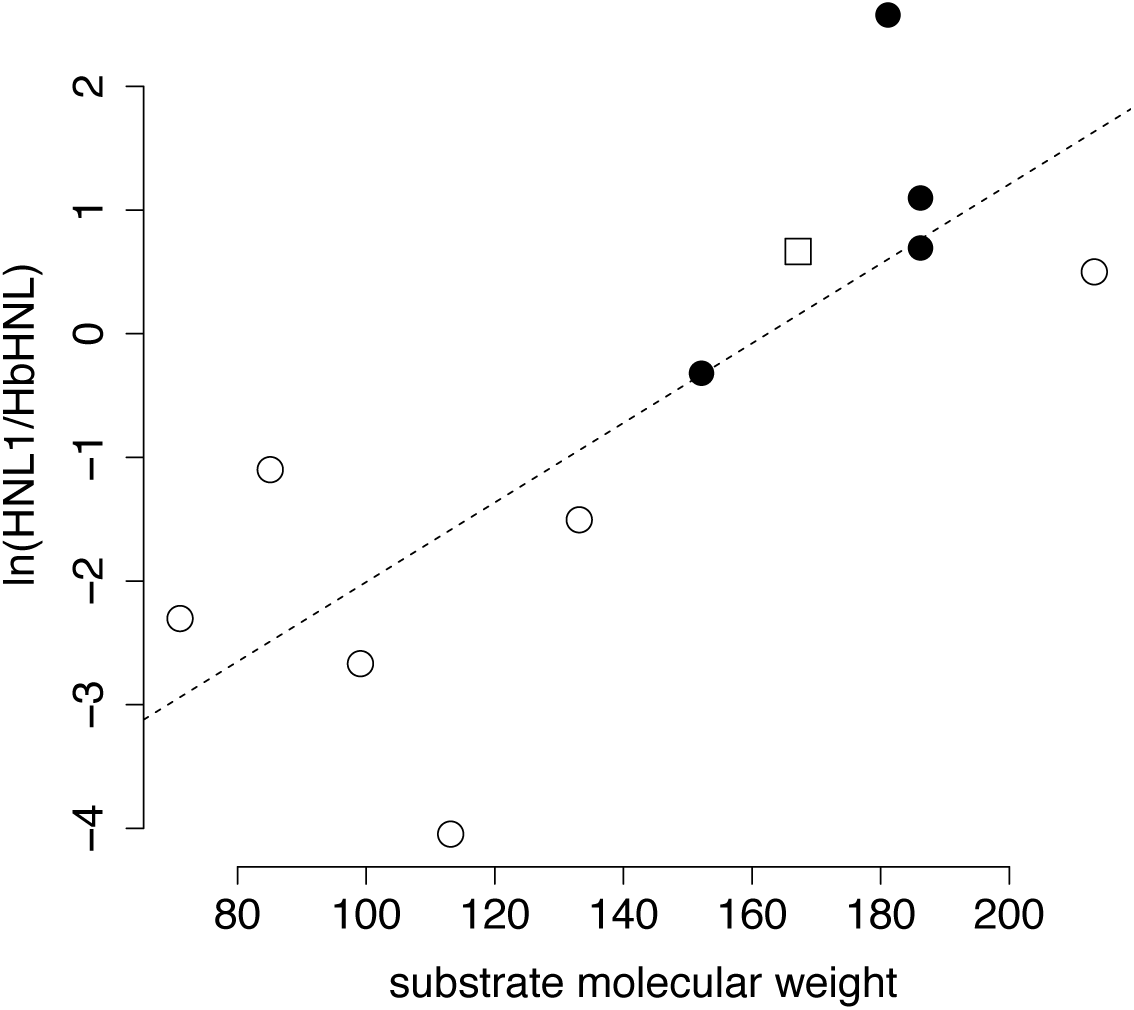
HNL1 favors larger substrates as compared to *Hb*HNL. The y-axis is the natural logarithm of the relative rate of the HNL1-catalyzed reaction versus the *Hb*HNL catalyzed reaction, while the x-axis is the molecular weight of the substrate, which approximates its size. The open symbols indicate lyase-type reactions: open circles are cleavage of cyanohydrins and the open square is the cleavage of 2-nitro-1-phenyl ethanol (reverse of a nitro-aldol addition). The filled black circles indicate hydrolysis of esters. The best linear fit (dashed line) is y = 0.032±0.009·x – 5±1 and the r^2^ = 0.61. Data for this figure are in S2 Table.

Catalytic promiscuity for hydroxynitrile lyases in this paper refers to their ability to catalyze ester hydrolysis. The comparison reactions are cleavage of mandelonitrile for the hydroxynitrile lyase activity and hydrolysis of *pNPAc* for the esterase activity. Both reactions can be measured colorimetrically. Both substrates contain one aromatic ring and have similar sizes (133 and 181 g/mol, respectively), but neither is a natural substrate. The natural substrate for *Hb*-HNL is acetone cyanohydrin [44], while the natural substrate for SABP2 is methyl salicylate [45]. We define this catalytic promiscuity as the ratio of the catalytic efficiency for the hydrolysis of *p*NPAc over the catalytic efficiency of cleavage of mandelonitrile, eq. 7.

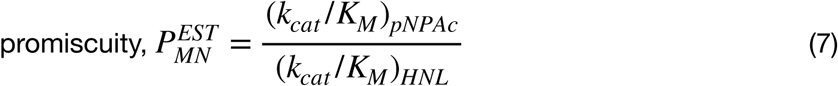

The two assays require slightly different pH, but are otherwise similar. Detecting the hydrolysis of *p*NPAc requires a pH above 7.2 so that the released the *p*-nitrophenol ionizes to the yellow *p*-nitrophenoxide. Cleavage of mandelonitrile requires a lower pH of 6.0 to minimize the spontaneous cleavage of mandelonitrile.

The ancestral enzyme HNL1 is approximately twice as promiscuous for esterase activity as compared to *Hb*HNL, Table 1. The catalytic efficiency (k_cat_/K_M_) of HNL1 is 2.5-fold higher than for *Hb*HNL for mandelonitrile cleavage, but is 4.5-fold higher for *p*NPAc hydrolysis. Thus, the promiscuity of HNL1 is two-fold higher that the promiscuity of *Hb*HNL. In both cases the promiscuous activity is ≥1000-fold slower than the cleavage of mandelonitrile. The site-directed mutagenesis data in Table 1 are described later in this paper. Other support for the promiscuity of HNL1 is its ability to catalyze the hydrolysis of naphthyl acetate esters, while no activity was detected with *Hb*HNL, S2 Table.

**Table 1.**
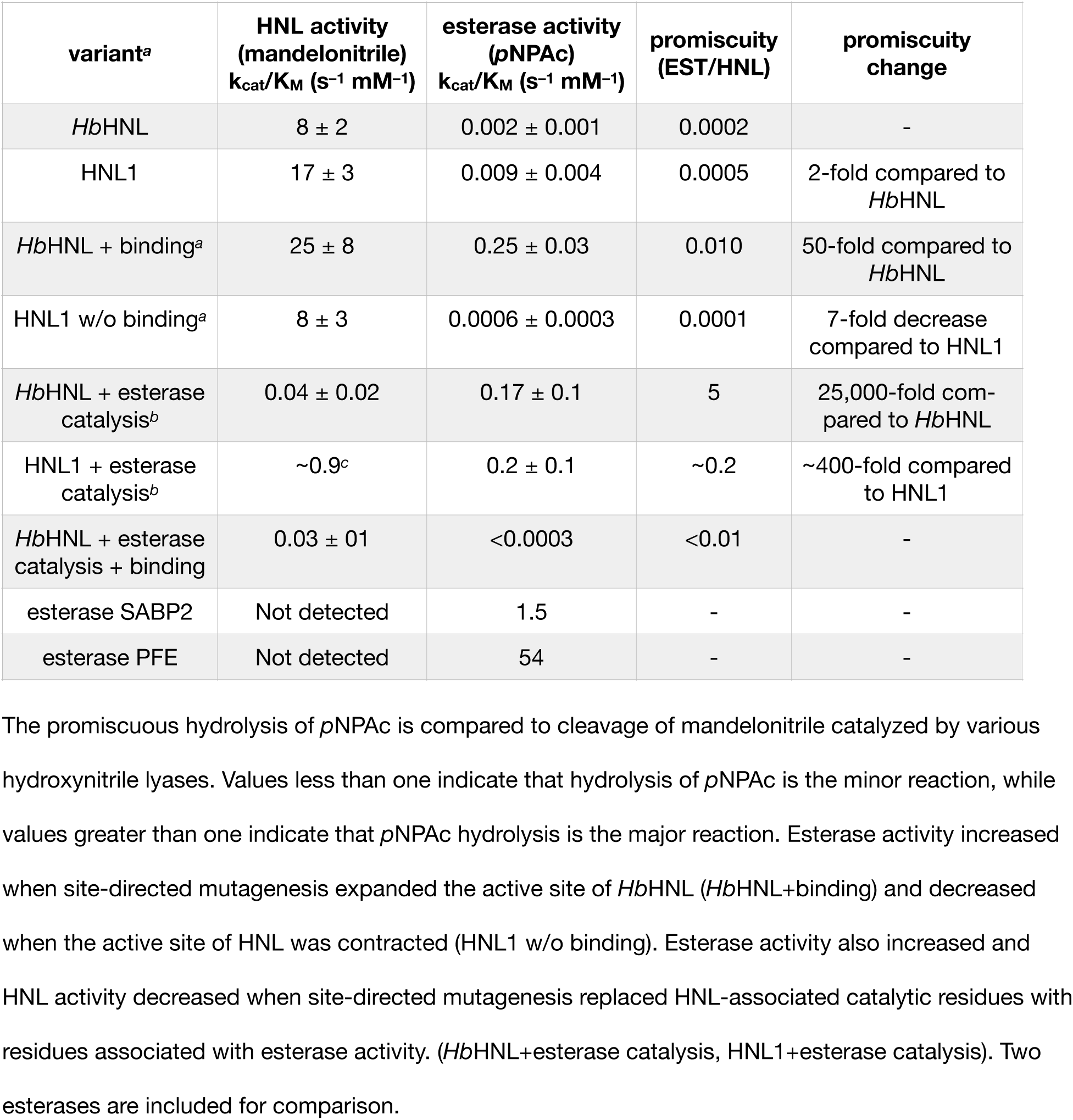
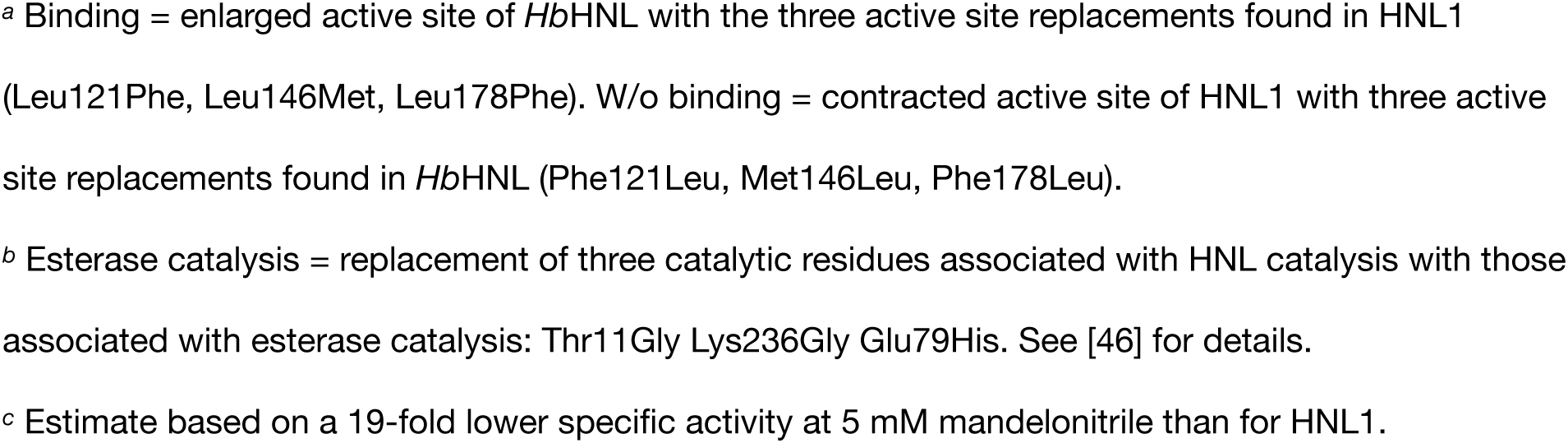
Catalytic promiscuity of hydroxynitrile lyases and variants.

| variant<sup>a</sup> | HNL activity<br>(mandelonitrile)<br>$k_{cat}/K_M$ ($s^{-1} mM^{-1}$) | esterase activity<br>(pNPAc)<br>$k_{cat}/K_M$ ($s^{-1} mM^{-1}$) | promiscuity<br>(EST/HNL) | promiscuity<br>change |
| --- | --- | --- | --- | --- |
| *HbHNL* | $8 \pm 2$ | $0.002 \pm 0.001$ | 0.0002 | - |
| HNL1 | $17 \pm 3$ | $0.009 \pm 0.004$ | 0.0005 | 2-fold compared to *HbHNL* |
| *HbHNL* + binding<sup>a</sup> | $25 \pm 8$ | $0.25 \pm 0.03$ | 0.010 | 50-fold compared to *HbHNL* |
| HNL1 w/o binding<sup>a</sup> | $8 \pm 3$ | $0.0006 \pm 0.0003$ | 0.0001 | 7-fold decrease compared to HNL1 |
| *HbHNL* + esterase catalysis<sup>b</sup> | $0.04 \pm 0.02$ | $0.17 \pm 0.1$ | 5 | 25,000-fold compared to *HbHNL* |
| HNL1 + esterase catalysis<sup>b</sup> | $\sim 0.9^c$ | $0.2 \pm 0.1$ | $\sim 0.2$ | $\sim 400$-fold compared to HNL1 |
| *HbHNL* + esterase catalysis + binding | $0.03 \pm 0.1$ | $<0.0003$ | $<0.01$ | - |
| esterase SABP2 | Not detected | 1.5 | - | - |
| esterase PFE | Not detected | 54 | - | - |

The promiscuous hydrolysis of pNPAc is compared to cleavage of mandelonitrile catalyzed by various hydroxynitrile lyases. Values less than one indicate that hydrolysis of pNPAc is the minor reaction, while values greater than one indicate that pNPAc hydrolysis is the major reaction. Esterase activity increased when site-directed mutagenesis expanded the active site of *HbHNL* (*HbHNL*+binding) and decreased when the active site of HNL was contracted (HNL1 w/o binding). Esterase activity also increased and HNL activity decreased when site-directed mutagenesis replaced HNL-associated catalytic residues with residues associated with esterase activity. (*HbHNL*+esterase catalysis, HNL1+esterase catalysis). Two esterases are included for comparison.
<sup>a</sup> Binding = enlarged active site of *HbHNL* with the three active site replacements found in HNL1
(Leu121Phe, Leu146Met, Leu178Phe). W/o binding = contracted active site of HNL1 with three active site replacements found in *HbHNL* (Phe121Leu, Met146Leu, Phe178Leu).
<sup>b</sup> Esterase catalysis = replacement of three catalytic residues associated with HNL catalysis with those associated with esterase catalysis: Thr11Gly Lys236Gly Glu79His. See [46] for details.
<sup>c</sup> Estimate based on a 19-fold lower specific activity at 5 mM mandelonitrile than for HNL1.

#### Thermostability of HNL1

Another property of HNL1 that differs from its closest modern descendent *Hb*HNL is thermostability. One way to measure thermostability is by comparing the midpoints of the transition between folded and unfolded structures as the proteins are heated. Since this transition is only partly reversible, it corresponds to an apparent, not true, melting temperature. The apparent melting temperature (T_m_) of HNL1 is 80-81 °C as measured by inherent fluorescence (S3 Fig). This melting temperature is 9 °C higher than for *Hb*HNL (T_m_ = 71 °C, S3 Fig) and about 10 °C higher than for *Me*HNL (∼70 °C) as measured by circular dichroism [47]. This 9-10 °C difference corresponds to an estimated 2.6 kcal/mol increase in the free energy of unfolding of HNL1 as compared to *Hb*HNL and *Me*HNL.

Besides a higher melting temperature, HNL1 is remarkably better at avoiding aggregation and irreversible inactivation. Upon heating for 15 min at 70 °C *Hb*HNL loses 50% of its activity. This stability varies slightly with buffer [48]. Similarly, *Me*HNL loses 50% of its activity upon heating for 30 min at 60 °C [47]. This inactivation stems from a fraction of the proteins unfolding at these temperatures below T_m_ and aggregating irreversibly. In sharp contrast, HNL1 requires heating to 95 °C, well above T_m_, for 30 min to inactivate 50% of its activity, S4 Fig. This 25-35 °C difference in inactivation temperature indicates that HNL1 effectively resists irreversible aggregation even when all the protein is unfolded.

Urea-induced unfolding measurements confirm that HNL1 is more stable than *Hb*HNL and suggest that unfolded HNL1 retains more folded structure than *Hb*HNL, Table 2 (S5 Fig shows experimental data). The concentration of urea required to unfold half of the proteins is higher for HNL1 (3.8±0.3 M) than for *Hb*HNL (2.9±0.1 M), which indicates that HNL1 is more stable than *Hb*HNL. The m-value for the urea unfolding experiment (absolute value of the slope of the plots in S5 Fig panel B), which indicates the sensitivity of the protein to urea and depends on the solvent accessible surface area exposed upon unfolding [49], is significantly higher for *Hb*-HNL (0.93±0.02 kcal/mol·M) than for HNL1 (0.61±0.03 kcal/mol·M). The m-value for *Hb*HNL is typical, but the m-value for HNL1 is unusually low. The x-ray structure below shows that the folded structures of both proteins are similar, so the lower m-value for HNL1 suggest that it exposes less solvent accessible surface area upon unfolding. The low sensitivity of HNL1 to unfolding by urea suggests that the unfolded form of HNL1 retains more folded structure than the unfolded form of *Hb*HNL. This retention of structure may account for the ability of HNL1 to refold and avoid aggregation upon heat unfolding. Robic et al. [50] found that ribonuclease H from thermophiles retained partial structure upon unfolding, but homologs from mesophiles unfolded more completely. Thermostable reconstructed ancestral ribonucleases also retained partially folded structure upon unfolding [51].

**Table 2.**
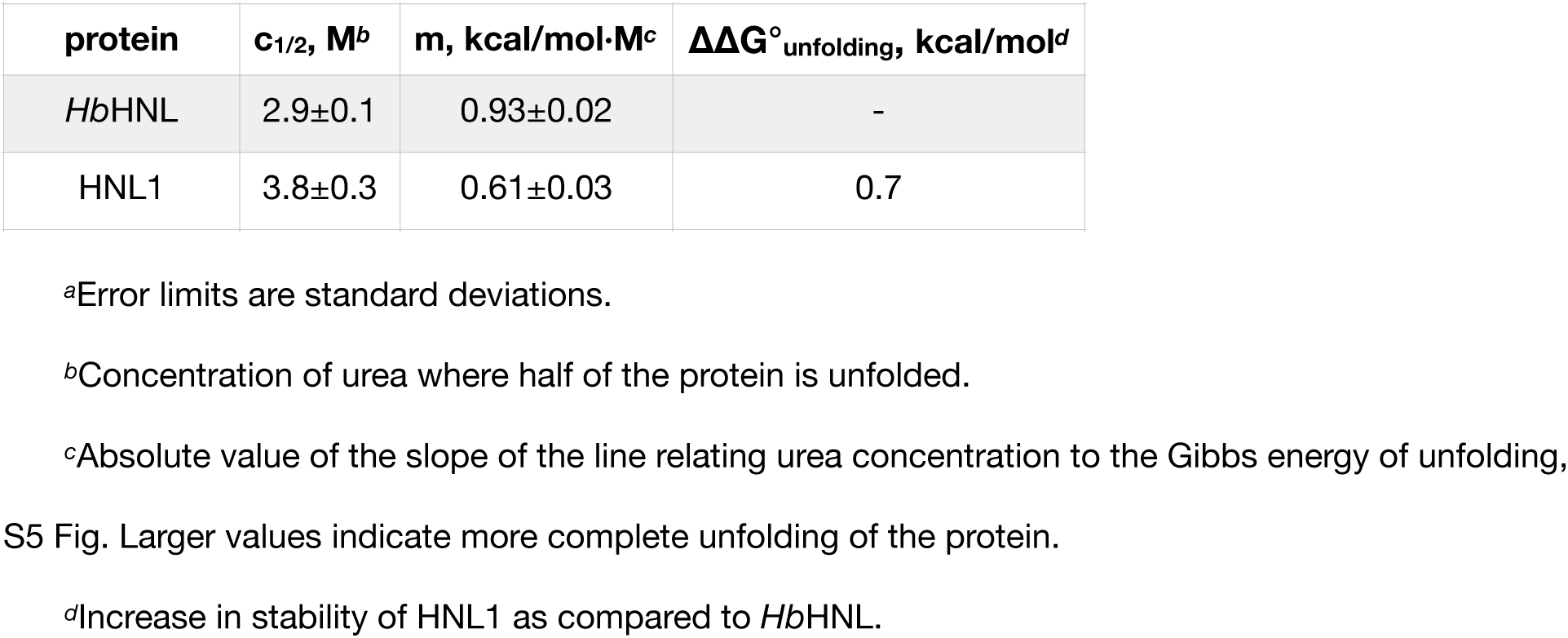
Urea-induced unfolding of *Hb*HNL and HNL1.^*a*^.

The Gibbs energy of unfolding for HNL1 is estimated to be 0.7 kcal/mol higher than for *Hb*-HNL. This increased stability is consistent with the 9 °C higher melting temperature of HNL1. When the slopes of the urea concentration versus unfolding Gibbs energy (m-values in Table 3) are similar, one can compare the extrapolated Gibbs energies of unfolding in pure water. This comparison incorrectly suggests that HNL1 is less stable than *Hb*HNL: ΔG°_H2O_ = 2.69±0.5 kcal/ mol for *Hb*HNL and 2.3±0.1 for HNL1. However, when these slopes differ, as they do in this case, the unfolding processes of the two proteins differs, so this comparison is misleading and suggests that HNL1 is less stable. To compare stability when the slopes differ, Pace and Scholtz [31] recommend comparing the urea concentrations at half unfolding. To convert these to Gibbs energies, one multiplies the difference in the half-unfolding-urea concentrations (0.9 M in this case) by the average of the slopes (0.72 kcal/mol·M), which yields 0.7 kcal/mol.

**Table 3.**
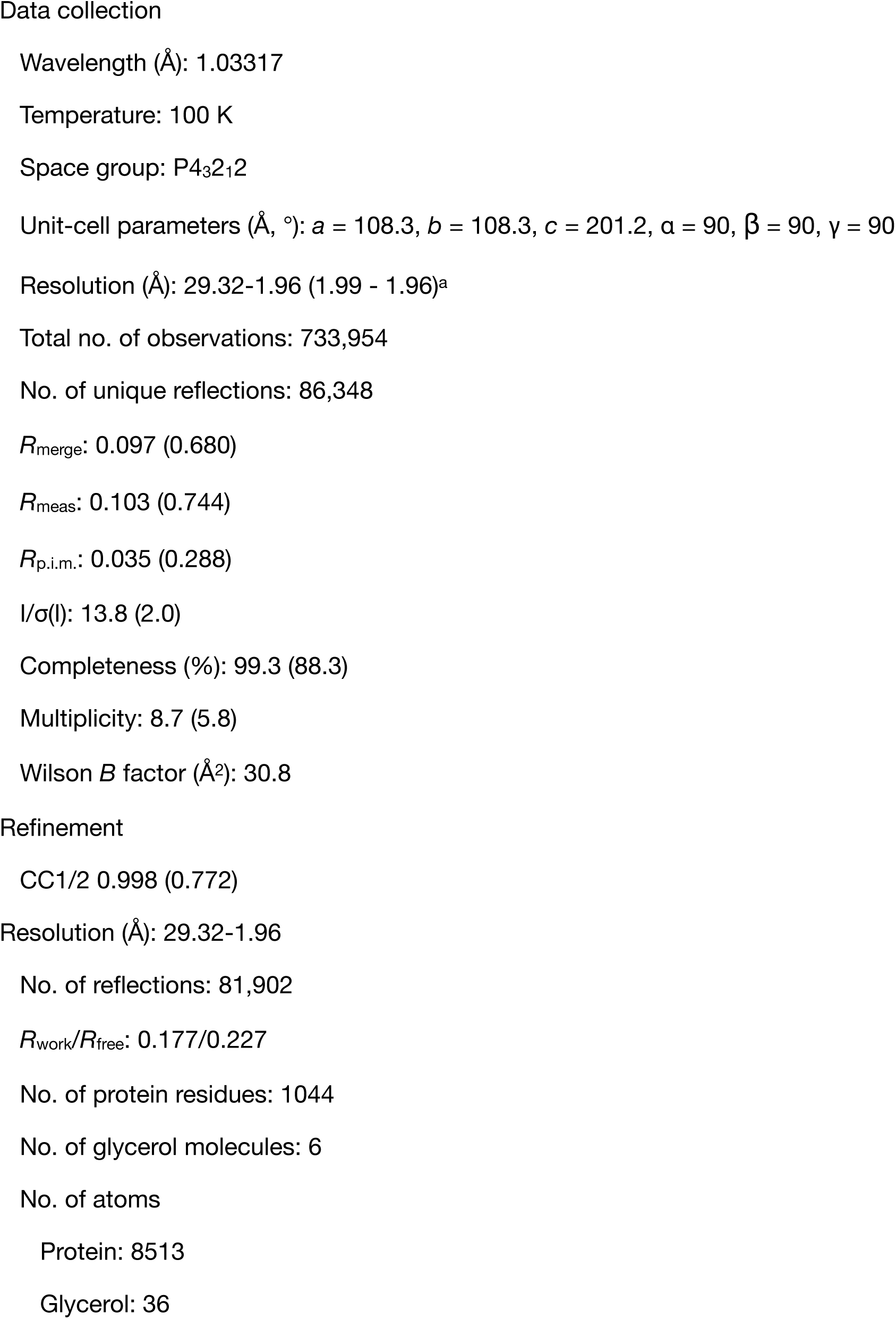
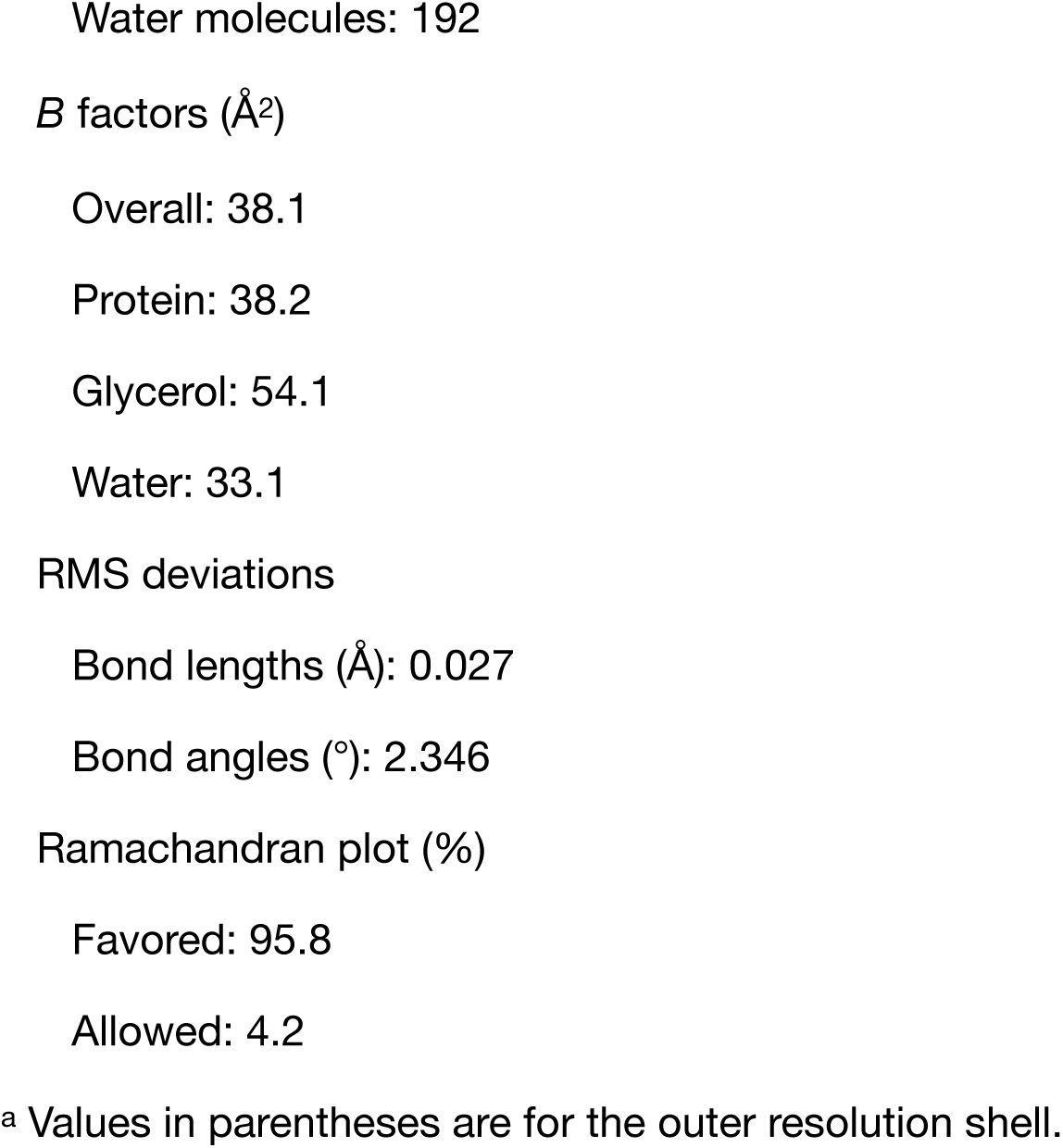
Data-collection and refinement statistics for HNL1 (pdb id 5tdx).

### X-ray structure of HNL1

We next solved the x-ray structure of HNL1 to identify structure differences that could explain the enhanced promiscuity. Octahedral crystals of HNL1 grew after 2-3 weeks in ammonium citrate and bis-tris propane solution at pH 6.8. The largest crystal - approximately 200 μm across - was harvested on day 20. This crystal diffracted well, and data were collected at beamline 23 IDB at the Advanced Photon Source, Argonne National Labs, by remote access. Molecular replacement using *Hb*HNL (pdb id 1yb6, 81% amino acid identity; 49 differing amino acids) found four monomers in the asymmetric unit. The refined model of HNL1 (pdb id 5tdx) contained a dimer of dimers. The four polypeptide chains; A, B, C, and D; comprised residues 2 through 261, 260, 265, and 261, respectively. The model contained six glycerol molecules, including one in each active site, and 192 water molecules. Crystallographic refinement resulted in a model to 1.96-Å resolution with an R_work_/R_free_ of 9.0/11.6%, and other statistics listed in Table 3. Electron density provided clear, unambiguous positioning of most residues including the active site residues, Fig 3.

**Fig 3.**
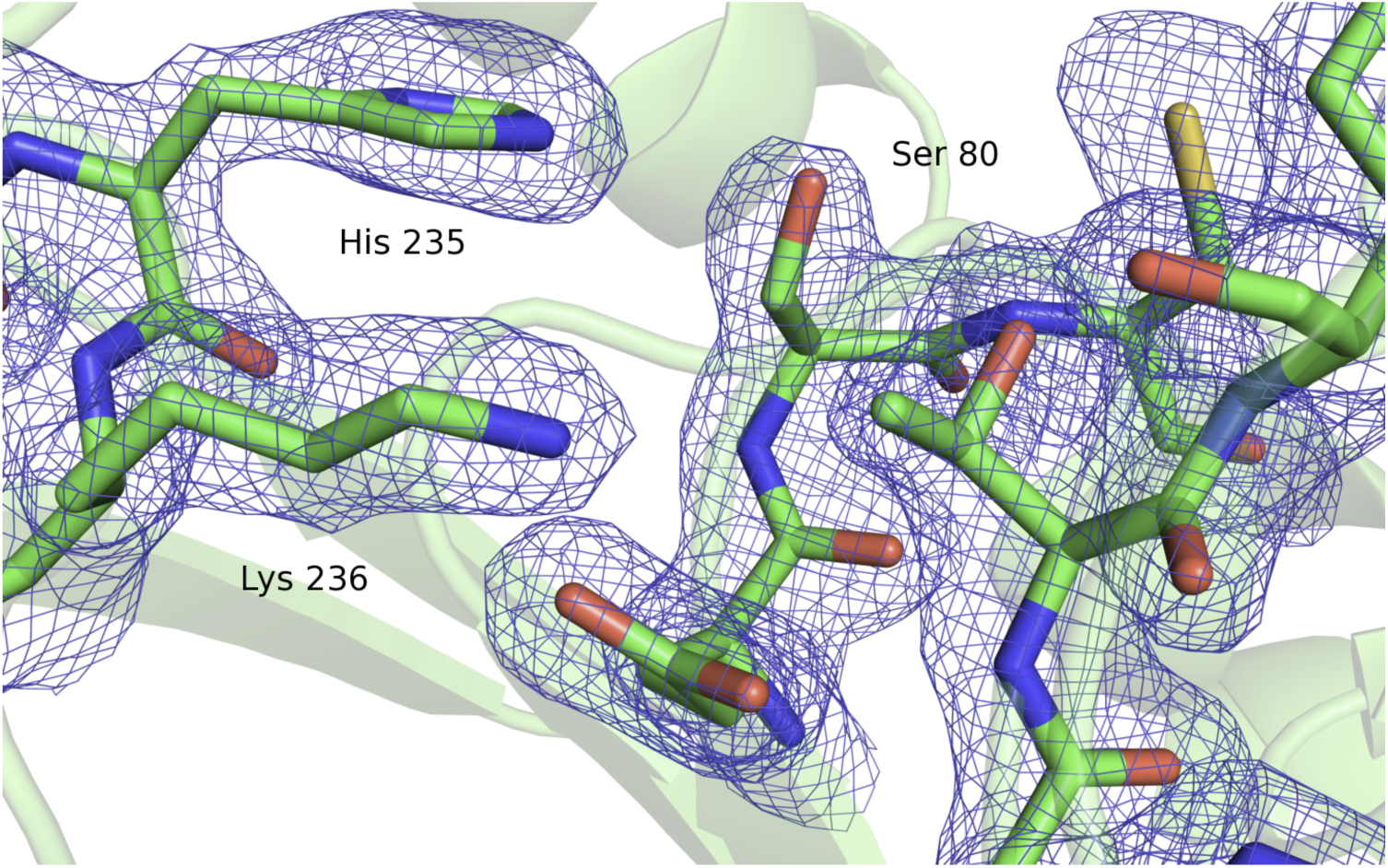
The active site of HNL1 showing electron density (2Fo − Fc) contoured at 1σ. The labels mark serine 80 and histidine 235 from the catalytic triad and the catalytically essential lysine 236.

The asymmetric unit contains a dimer of dimers. Chains A and B form one dimer, Fig 4; chains C and D form the second dimer, and the interaction of these two dimers appears to result from crystal packing. Modern hydroxynitrile lyases form dimers both in crystals and in solution [52-56]. For HNL1, the calculated binding between chains A and B is strong (−16.8 kcal/ mol) and statistically significant (p-value = 0.013) according to the PISA assessment of macromolecular interfaces [57]. The calculated binding between the dimers is weak and not statistically significant confirming that this interaction is the result of crystallization. Therefore, the native arrangement of HNL1 is as the dimer like the modern hydroxynitrile lyases discussed here.

**Fig 4.**
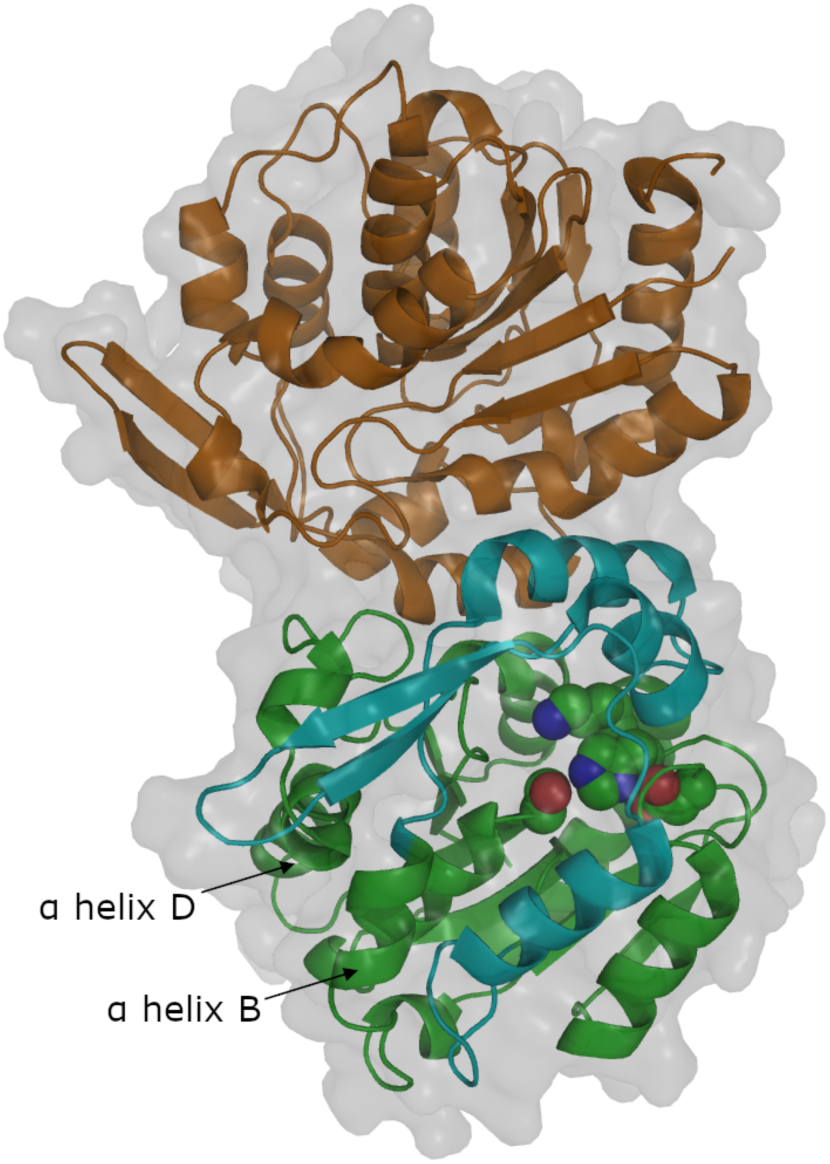
Chains A and B form a dimer in HNL1. The green cartoon indicates the core domain of chain A and the cyan cartoon indicates the cap domain of chain A. The catalytic triad (Ser80-His235-Asp207) and the catalytic lysine 236 are shown as space-filling. The orange cartoon indicates chain B of the dimer. The white surface indicates the solvent accessible surface. The labelled α-helices are discussed later in the paper.

Like other α/β-hydrolase-fold enzymes, the HNL1 monomer consists of two domains: a catalytic and a cap (or lid) domain, Fig 4. The catalytic domain consists of residues 1-107 and 180-265 and contains the catalytic triad (Ser 80, His 235, and Asp 207) as well as the catalytic Lys 236. The cap domain consists of residues 108-179 and forms part of the substrate-binding portion of the active site. The catalytic domain includes a core of six parallel beta strands which are surrounded by alpha helices as is typical of α/β-hydrolase-fold enzymes. Helices and strands alternate sequentially within the catalytic domain, S6 Fig. All proline residues adopt the trans conformation along the peptide bond.

The 49 amino acids that differ between HNL1 and *Hb*HNL occur throughout the protein, but occur more frequently in the lid domain and on the surface of the protein, S3 Table, S7 Fig. They are also more likely to be non-conservative replacements in these regions. These larger changes are consistent with more frequent changes during evolution in the lid domain as compared to the catalytic domain and on the protein surface as compared to the interior of the protein. The fraction of replacements in the lid domain (25%) is larger and more likely to be non-conservative (66.7% of the replacements) than in the catalytic domain: 16.8% replacements, 32.3% non-conservative. Overall about half of the replacements are conservative replacements (27, 55% of 49), while the remaining 22 are non-conservative replacements. Most of the residues in HNL1 (188 of 257, 73.2%) have at least some of their surface exposed to the solvent, S3 Table. The replaced residues skew even more strongly to the surface: 40 of the 49 replacements, 81.6%. All but one of the non-conservative replacements occur on the surface.

The overall structure of HNL1 is similar to that of modern hydroxynitrile lyases *Hb*HNL, *Me*HNL, and *Bm*HNL. The RMSD between Cα’s (chain A in all cases) is 0.40 Å for *Hb*HNL (pdb id 3c6x, comparison of 217 amino acids), 0.47 Å for MeHNL (pdb id 1eb9, comparison of 236 amino acids), and 0.45 Å for *Bm*HNL (pdb id 3wwp, comparison of 220 amino acids). Both the catalytic triad (Ser 80, His 235, and Asp 207) as well as the catalytic Lys236, have the same orientations in the ancestral HNL1 and the modern HNLs. The structures are similar despite the 49 amino acid changes (19%) between HNL1 and *Hb*HNL, including 22 that are not chemically similar. The similarity in secondary, tertiary, and quaternary structure despite the numerous changes to the primary structure is consistent with the observed catalytic activity of HNL1. Taken together, the structure and properties of HNL1 are consistent with an ancestor of modern hydroxynitrile lyases.

#### Substrate-binding site differences

The substrate-binding site is defined as those residues that may directly contact the substrate mandelonitrile. There are 21 amino acid residues with at least two atoms within 7 Å of the bound mandelonitrile substrate in the x-ray structure of *Hb*HNL (pdb id 1yb6). Among these 21, there are three substitutions in HNL1: Leu121, Leu146 and Leu178 in *Hb*HNL are replaced by Phe121, Met146 and Phe178 in HNL1, S6 Fig. Although the Phe and Met residues are longer than Leu and penetrate further into the substrate binding cavity, Fig 5, the active site volume of HNL1 is larger than the active site volume of *Hb*HNL. The larger phenylalanines in HNL1 push the side chain of Trp128 out of the active site by 1.3 Å as compared to *Hb*HNL (1 Å relative to *Me*HNL), thereby creating a larger substrate-binding region in HNL1. An overlay of the surface of the substrate-binding region of HNL1 and *Hb*HNL, Fig 5, shows the differences in shape of these regions. While the larger amino acids in HNL1 cause parts of the substrate binding pocket to be smaller, HNL1 provides a 1.3 Å longer pocket at the for the aromatic ring of mandelonitrile based on the *Hb*HNL complex.

**Fig 5.**
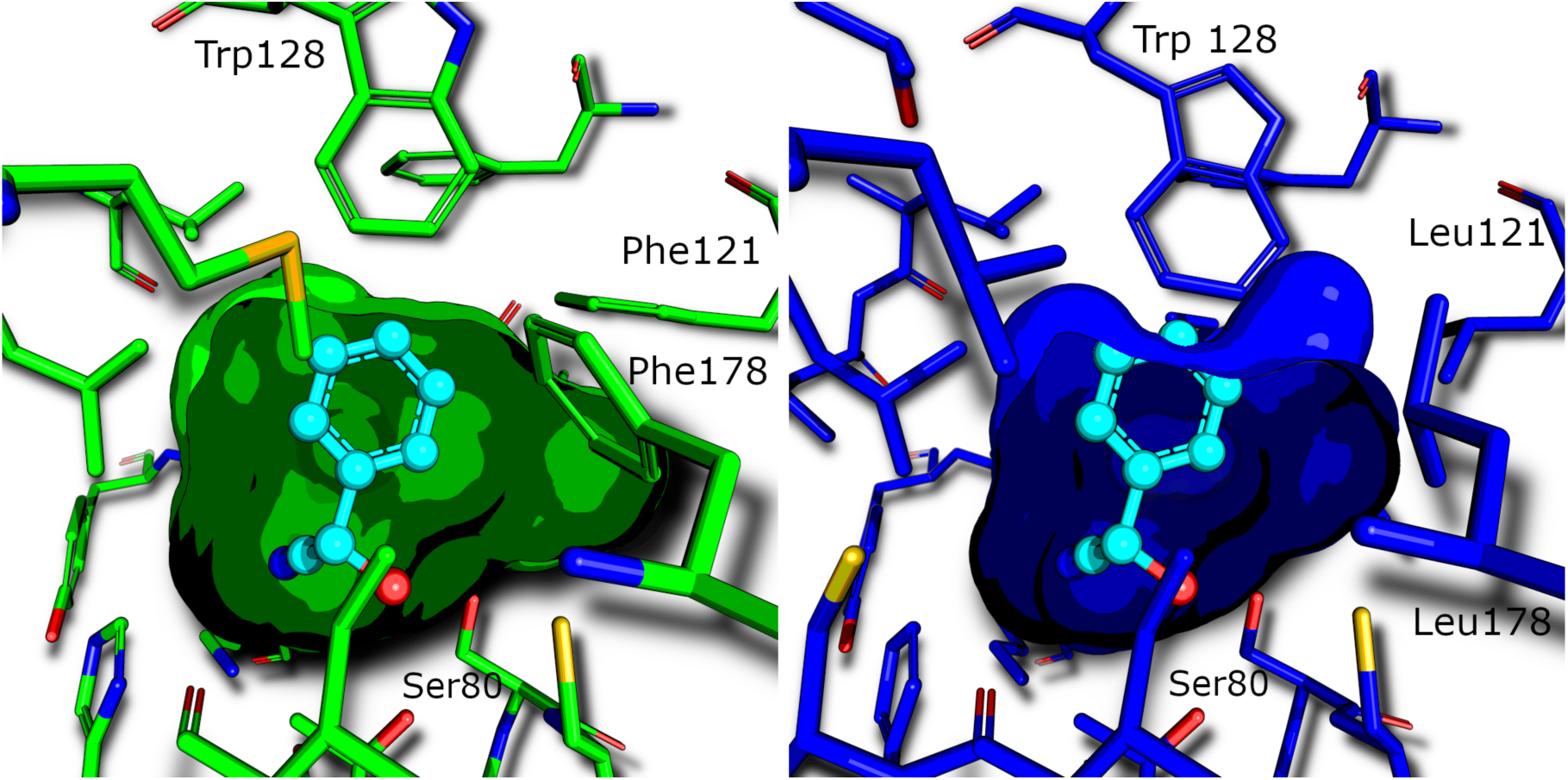
The active site of HNL1 (pdb id 5tdx, green) is larger than the active site of *Hb*-HNL (pdb id 3c6x, blue). Since the structure of HNL1 does not contain bound mandelonitrile, the structure of *Hb*HNL chosen for comparison also lacks bound mandelonitrile. However, mandelonitrile (cyan balls and sticks) has been added as found in another structure of *Hb*HNL (pdb id 1yb6) to orient the viewer. The phenylalanines at positions 121 and 178 in HNL1 push Trp128 away from the active site. The closest distance between Ser80 and Trp128 is 8.2 Å in HNL1, but only 6.9 Å in *Hb*HNL. S8 Fig shows a overlay of HNL1 and *Hb*HNL active sites in stereo view.

The active site volume of HNL1 (143 Å^3^) is larger than the active site volumes of *Hb*HNL (94-119 Å^3^) and esterase SABP2 (123 Å^3^). These active sites were defined as the innermost section of the tunnel up to the first pinch point, which is a region where the tunnel narrows, then widens again. The start of the tunnel *Hb*HNL containing bound mandelonitrile (pdb id 1yb6) was used as the starting point of the active site for all structures. The end of of the active site was defined as 10.5 Å from the starting point for all structures. The tunnel for *Hb*HNL containing bound mandelonitrile (pdb id 1yb6) showed a pinch point at 10.3 Å and *Hb*HNL without bound substrate (pdb id 3c6x) showed a pinch point at 10.7 Å. The tunnel for HNL1 narrowed at ∼10.5 Å, without a clear pinch point. The tunnel for SABP2 showed no narrowing or pinch point. To make a fair comparison, the end of the active site was defined as 10.5 Å for all enzymes, S10 Fig, S11 Fig. The volumes of the active sites were calculated using MOLEonline 2.0 [42] using default settings.

Another size comparison is the distance between the catalytic Ser80 and Trp128, Table 4. The substrate binds between these two residues, so the distance between them limits substrate size and orientation. By this distance measure, the substrate-binding site of HNL1 is also larger (8.2 Å) than that of *Hb*HNL (6.9 Å) and *Me*HNL (7.2 Å). The binding sites in *Bm*HNL and esterase SABP2 are nearly as large as HNL1. Trp128 in *Bm*HNL is in a similar position as in HNL1 [58], despite lower overall sequence similarity (68% identity) than to *Hb*HNL (81%) and *Me*HNL (77%). Like HNL1, *Bm*HNL accepts large aromatic substrates [53], but BmHNL’s natural substrate is unknown. The position of the tryptophan is also similar in the related esterase SABP2 from *N. tabacum* [59]. Esterase PNAE differs from SABP2 since the smaller amino acid methionine replaces Trp128. This replacement creates a substrate binding region even larger than observed in HNL1.

**Table 4.**
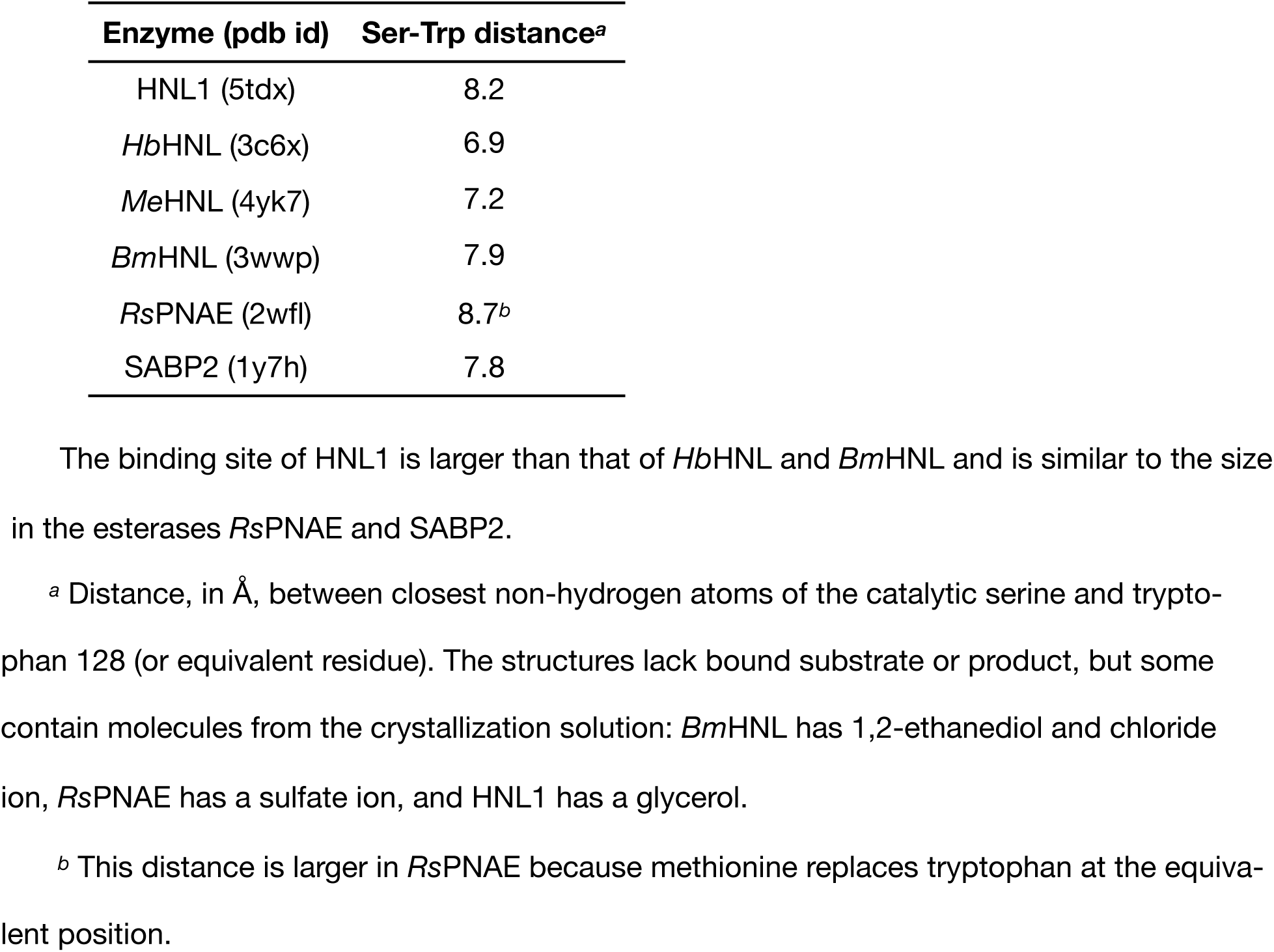
Size of the substrate-binding region as measured by the distance between the catalytic serine and Trp128 (or equivalent).

| Enzyme (pdb id) | Ser-Trp distance<sup>a</sup> |
| --- | --- |
| HNL1 (5tdx) | 8.2 |
| *HbHNL* (3c6x) | 6.9 |
| *MeHNL* (4yk7) | 7.2 |
| *BmHNL* (3wwp) | 7.9 |
| *RsPNAE* (2wfl) | 8.7<sup>b</sup> |
| SABP2 (1y7h) | 7.8 |

The binding site of HNL1 is larger than that of *HbHNL* and *BmHNL* and is similar to the size in the esterases *RsPNAE* and SABP2.
<sup>a</sup> Distance, in Å, between closest non-hydrogen atoms of the catalytic serine and tryptophan 128 (or equivalent residue). The structures lack bound substrate or product, but some contain molecules from the crystallization solution: *BmHNL* has 1,2-ethanediol and chloride ion, *RspNAE* has a sulfate ion, and *HNL1* has a glycerol.
<sup>b</sup> This distance is larger in *RspNAE* because methionine replaces tryptophan at the equivalent position.

In addition to a larger substrate-binding region in HNL1 as compared to *Hb*HNL, this substrate-binding region is more dynamic in HNL1. Comparison of the normalized crystallographic B-factors of HNL1 with those for *Hb*HNL reveal increased dynamics in the substrate-binding region of HNL1, Fig 6 and S12 Fig. Crystallographic B-factors indicate the uncertainty in atom position due to thermal motion (dynamics) and due to disorder (defects in crystal packing). B-factors also vary with the resolution of the structure and with crystal packing, so they may not indicate the flexibility in solution, but it can still be informative to make these comparisons. Comparison of the B-factors for the Cα atoms reveals differences in main chain flexibility and ignores differences in side chain movements. To compare different structures, the B-factors of each residue was normalized to the mean B-factor of the Cα atoms. Both structures had similar resolution: 1.96 Å for HNL1 (5tdx) and 1.8 Å for *Hb*HNL (1sc9). Several regions in HNL1 show increased flexibility as compared to *Hb*HNL. The most relevant to catalytic promiscuity are three regions with increased flexibility that shape the active site. These regions are from slightly more than one to over three standard deviations more flexible than the corresponding residues in *Hb*HNL. This higher flexibility may create alternative conformations that enable promiscuous ester hydrolysis.

**Fig 6.**
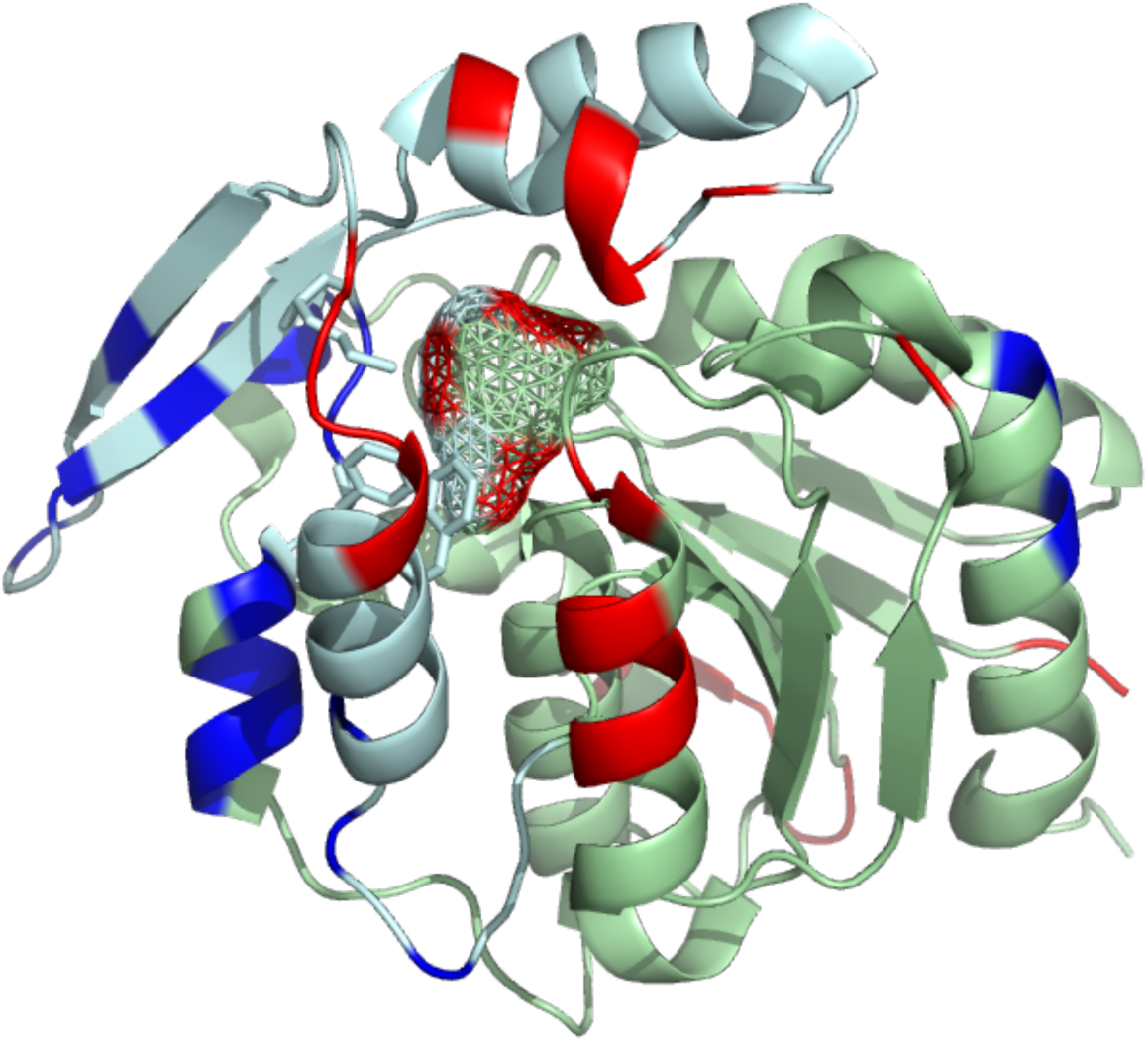
Three regions of the substrate-binding site of HNL1 are more flexible (red) than the corresponding regions of *Hb*HNL. The flexibility differences between HNL1 (pdb id 5tdx, chain A) and *Hb*HNL (pdb id 1sc9) are mapped onto a cartoon representation of the HNL1 structure as indicated by differences (HNL1 minus *Hb*HNL) in the normalized B-factors. The light blue cartoon representation indicates the lid domain, while the light green indicates the catalytic domain. Amino acids whose Cα atoms are at least one standard deviation more flexible in HNL1 than in *Hb*HNL are colored red, while less mobile amino acids are blue. The side chains of the three amino acids (Phe121, Met146, Phe178) responsible for the larger active site in HNL1 are shown as sticks. The mesh shows the space available for substrate binding. The projection of red color onto three regions of this mesh indicates that these three regions of the substrate-binding site are more flexible in HNL1 than in *Hb*HNL.

Docking of *pNPAc* into the active site of HNL1 suggests that the larger volume allows the substrate to explore a wider range of orientations. For catalysis, the *pNPAc* must orient with the acetate group and *p*-nitrophenyl groups in the acyl and leaving pockets, respectively.

Docking (SwissDock) identified 255 potential orientations of *pNPAc* in the active site of each enzyme and grouped similar orientations into clusters. Three of the top ten docking clusters oriented *pNPAc* into the correct pockets for *Hb*HNL and HNL1, but the catalytically non-competent orientations differed between the two enzymes. For *Hb*HNL the major incorrect orientation exchanged locations of the acetate and *p*-nitrophenyl groups making catalysis impossible. For HNL1, four of the incorrect orientations were sideways orientations within the pocket that likely could rotate into a correct orientation for catalysis. This ability to explore new substrate binding orientations in HNL1 due to the larger active site may contribute to its higher promiscuous esterase activity as compared to *Hb*HNL.

#### Structure differences related to thermostability

The numbers of intra-protein interactions in HNL1 and *Hb*HNL are similar and do not suggest that their stabilities would significantly differ, S4 Table. The total solvent accessible surface area of HNL1 is 1.2% larger than *Hb*HNL. Since the structure of HNL1 contains 1.5% more amino acids residues in each monomer (261 vs. 257 for *Hb*HNL), this difference suggests similar compactness for both structures. On one hand, HNL1 shows slightly more hydrophobic interactions: a 2.3% higher number of non-polar contacts and 1.5% less non-polar surface area at the surface. On the other hand, *Hb*HNL shows slightly more electrostatic interactions: a 1.8% higher number of oppositely charged atoms within 7 Å of each other (111 vs 109) and a 2.1% higher number of hydrogen bonds (143 vs. 140).

Although the numbers of interactions are similar, an examination of individual interactions shows at least one hydrogen bond network that is stronger in HNL1 than in *Hb*HNL, Fig 7. In *Hb*HNL, this hydrogen bond network incorporates bridging water molecules, while in HNL1, the hydrogen bond network involved direct hydrogen bonds between protein atoms. The direct interaction is expected to be more favorable since it does not require the immobilization of water molecules.

**Fig 7.**
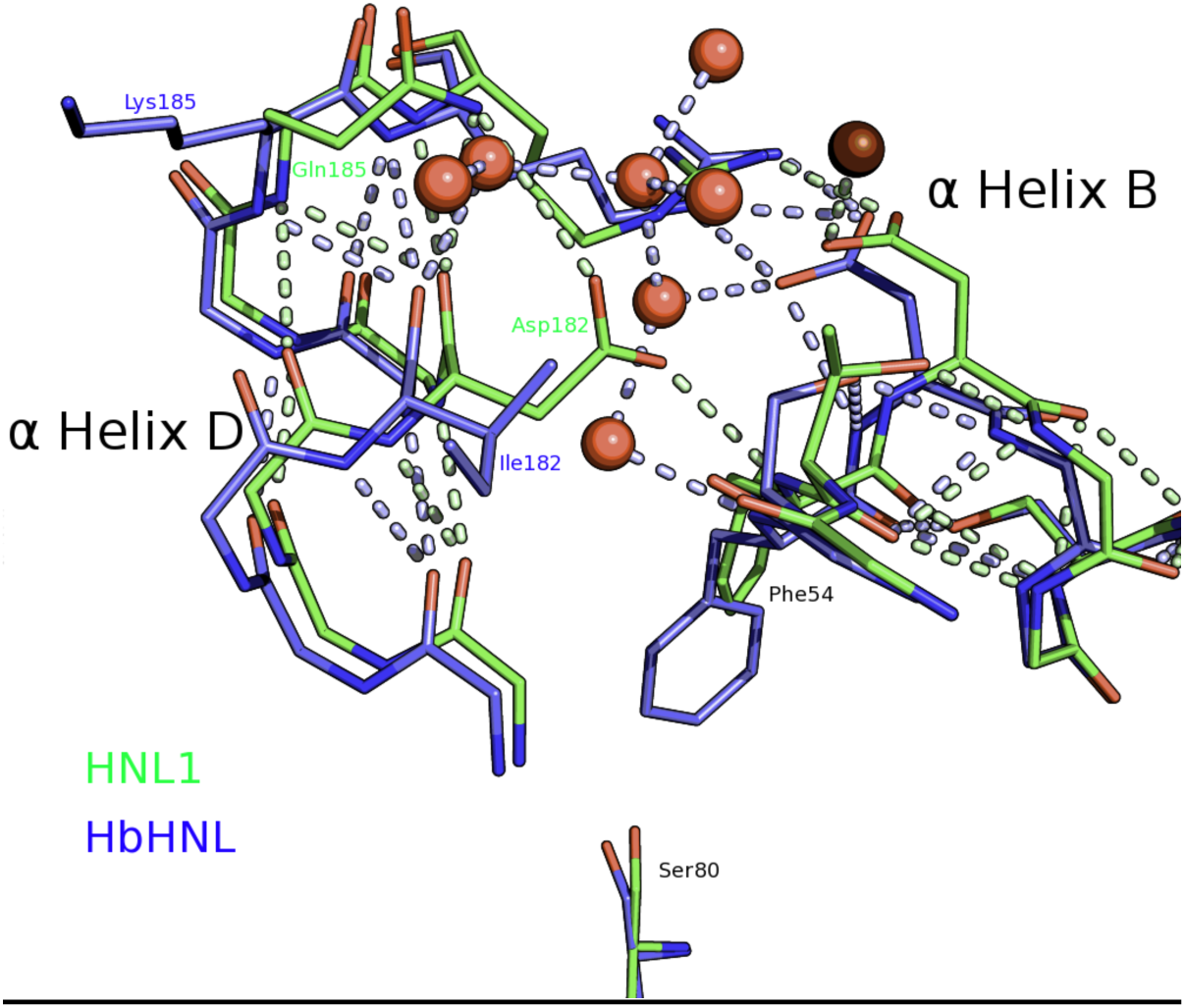
Hydrogen bonding network in HNL1 may contribute to higher thermostability. Aspartic acid 182 in HNL1 (green) hydrogen bonds directly to glutamine 185 and the backbone nitrogen of phenylalanine 54. This interaction bridges α helix B and the small α helix D. In contrast, *Hb*HNL (blue) hydrogen bonds indirectly to these helices via several water molecules (red spheres) that are absent from HNL1 (dark red sphere). Dashed lines indicate polar contacts. The bottom of the figure shows the catalytic serine for reference. The helices lie on the surface of the catalytic domain, but are not part of the substrate binding site. Fig 4 above shows the location of these helices within the whole structure.

HNL1 contains three fewer glycine residues than *Hb*HNL (14 vs 17; ignoring a glycine in the linker to the histidine tag) and one more proline residue (13 vs 12). These differences are consistent with a stabilization of HNL1 toward unfolding by reducing the flexibility of the unfolded form. Glycine is the most flexible amino acid residue, so replacing glycines with residues having fewer degrees of freedom reduces the flexibility of the unfolded form and shifts the balance toward the folded form. Proline is the least flexible amino acid residue, so replacing any residue with proline similarly reduces the flexibility of the unfolded form and shifts the balance toward the folded form.

One possible reason for the ability of HNL1 to resist aggregation is the lack of aggregation-prone regions, but Tango calculations rule out this explanation. The Tango algorithm identifies hydrophobic, β-sheet forming regions as possible sites for nucleation of aggregation [60]. Tango predicted several slightly aggregation prone regions in *Hb*HNL and *Me*HNL and one strongly aggregation-prone region in HNL1 (residues 200-204: VYVWT, corresponds to strand β5), which is not consistent with the observation that HNL1 is less prone to nucleation (S13 Fig).

### Site-directed mutagenesis

To test whether the enlarged binding site causes the increased promiscuity, we used site-directed mutagenesis to add or remove the three residues associated with the larger binding pocket, Table 1. First, we replaced these three residues in *Hb*HNL with the corresponding ones in HNL1 to expand its binding site (*Hb*HNL + binding, Table 2; Leu121Phe Leu146Met Leu178Phe). This expansion in *Hb*HNL increased the esterase activity 125 fold. This replacement also increased HNL activity 3 fold, so the promiscuity increased 50 fold. This promiscuity is 20-fold higher than for the ancestral enzyme HNL1. This large increase in promiscuity, caused by the large increase in esterase activity, is consistent with the notion that the larger active site increases promiscuous esterase activity. In a complementary experiment, we contracted the active site of HNL1 by replacing the three residues with the corresponding residues in *Hb*HNL (HNL1 w/o binding). The esterase activity of HNL1 w/o binding decreased 15 fold as compared to HNL1 and the HNL activity decreased 2 fold. The promiscuity of HNL1 w/o binding decreased 7 fold as compared to HNL1, mainly due to the decrease in esterase activity. This change is also consistent with the notion that reducing the size of the binding site would decrease esterase activity.

Expanding the binding site was not as effective in introducing esterase activity as the introduction of catalytic residues associated with esterase activity. In previous work, we switched the HNL activity of *Hb*HNL to esterase activity by replacing the catalytic residues associated with HNL activity in *Hb*HNL with the corresponding residues in esterases (Thr11Gly, Lys236Gly, Glu79His; [46]. This replacement decreased HNL catalytic efficiency 200 fold and increased esterase activity 85 fold. The major activity switched from HNL activity to esterase activity, but the catalytic efficiency was low compared to natural esterases. Here, we tested this same replacement in HNL1 (Thr11Gly Lys236Gly Glu79His = esterase catalysis), hypothesizing that the ancestral enzyme would be more malleable and result in a more efficient esterase. The HNL catalytic efficiency decreased 10-fold while the esterase catalytic efficiency increased 22 fold. This HNL1+esterase catalysis variant was similarly effective as an esterase, but retained more HNL activity. The esterase activities of *Hb*HNL+esterase catalysis and HNL1+esterase catalysis were ∼10-fold lower than that for a related esterase (SABP2).

An attempt to increase the esterase activity of *Hb*HNL+esterase catalysis by expanding its binding site was unsuccessful. The resulting enzyme (*Hb*HNL+esterase catalysis+binding) showed no detectable esterase activity and a drop in HNL activity compared to *Hb*HNL+esterase catalysis, Table 1. This lack of success indicates incompatibility between the two sets of substitutions, which individually increased esterase activity.

## Discussion

The three previously identified mechanisms for catalytic promiscuity are 1) similar transition states, 2) different active site conformations, and 3) rich catalytic network within the active site. The transitions states and mechanisms for ester hydrolysis and cyanohydrin cleavage differ significantly. There is some evidence for differences in active site conformations since the x-ray structure suggests increased flexibility of the residues in the substrate-binding site of HNL1 as compared to *Hb*HNL. These differences are likely small local changes, not large shifts in secondary structure elements. The rich catalytic network within the active site is likely the most important for catalytic promiscuity of HNL’s. The active site serine can act as a nucleophile for ester hydrolysis and or as a base for cyanohydrin cleavage, Fig 9. But exploiting this rich catalytic network also requires different orientations of the substrate. The three-fold higher catalytic promiscuity of ancestral hydroxynitrile lyase HNL1 and compared to modern *Hb*HNL is likely due to a larger substrate binding site in HNL1 compared to *Hb*HNL. This larger substrate-binding site allows the promiscuous substrate, *p*NPAc, to orient in a catalytically productive manner. The comparison substrates, mandelonitrile and *p*NPAc, both contain aryl groups, but the catalytically productive orientation of these aryl groups differs, Fig 8. If one divides the active site into an acyl-binding region and an alcohol-binding region, then the productive orientation of *p*NPAc places the aryl group in the alcohol-binding region, but the productive orientation of mandelonitrile places the aryl group in the acyl-binding region. This differing requirement for the substrate orientation creates the expectation that a large active site that could accommodate both orientations would lead to increased catalytic promiscuity. The x-ray structure shows a larger active site in HNL1 as compared to *Hb*HNL, so this difference can contribute to the 3 fold higher promiscuity of HNL1 for ester hydrolysis. Higher catalytic promiscuity in the alkaline phosphatase family was also associated with a larger active site [11].

**Fig 8.**
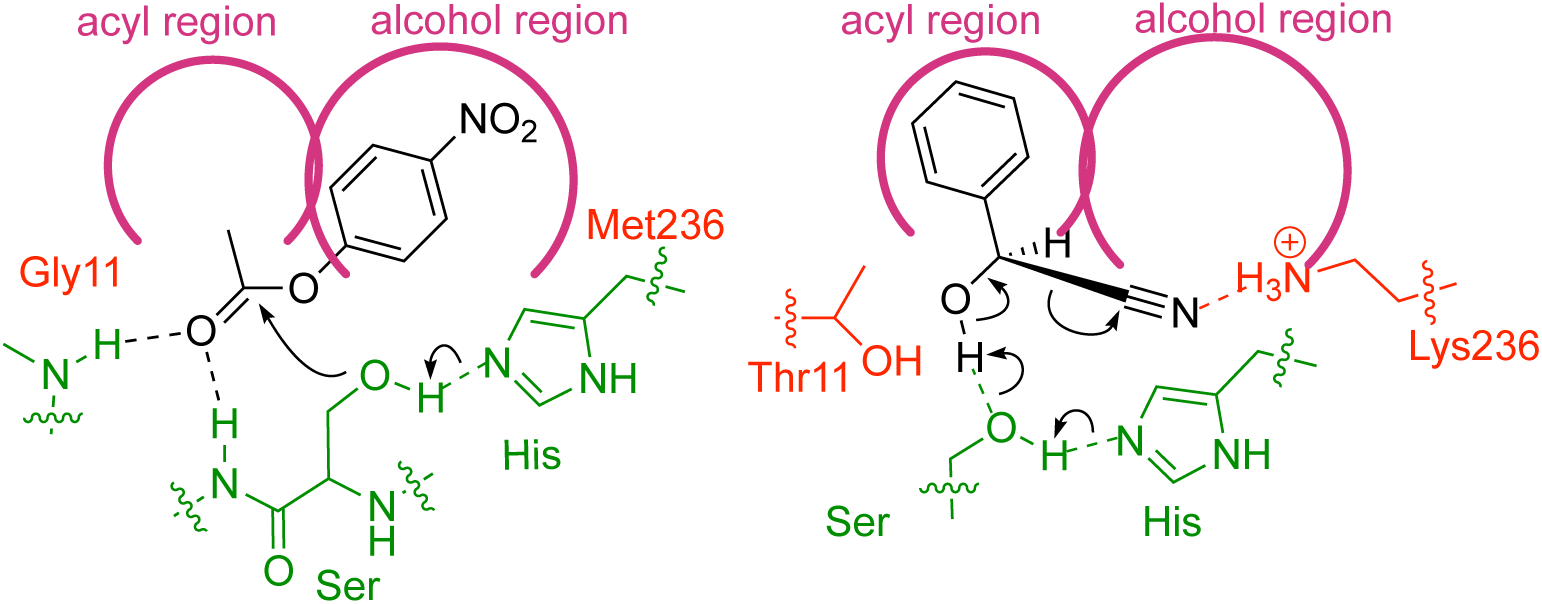
The aryl group placement differs for *p*NPAc and mandelonitrile. During ester hydrolysis (left scheme), the *p*-nitrophenyl moiety of the ester orients in the alcohol region of the substrate binding site. During cyanohydrin cleavage, the phenyl group of mandelonitrile orients in the acyl region of the substrate-binding site. The catalytic triad and oxyanion hole residues are in green, the residues associate with ester catalysis (Gly11, Met236) or HNL catalysis (Thr11, Lys236) are shown in red, the substrates are in black and the substrate binding site is shown schematically in magenta.

One proposed reason for the higher thermostability of ancestral proteins is that the average earth temperatures were higher in the past [14-18, 61]. Adaptation to a warmer environment could only account for part of the 9-10 °C increase in melting temperature of HNL1. Plant defense enzymes like HNL1 likely formed during the Cretaceous period approximately 100 million years ago as flowering plants and insects co-evolved. Modern descendants of HNL1 belong to the Euphorbiaceae family of flowering plants, which diverged between 110 and 69 million years ago [62, 63]. Global temperatures during this period were only ∼4 °C warmer than today [64]. Thermostability may also be a byproduct of stability to other environmental stresses such as high levels of oxidative stress or radiation [65].

A second reason for high thermostability of resurrected ancestral proteins is reconstruction bias toward stabilizing substitutions [65-67]. Stabilizing residues are more likely to be conserved in modern descendants, and thus the identity of these residues is available to correctly infer the ancestral residue, while destabilizing residues are more likely to be lost in the descendants and therefore are more likely to be missed while inferring the ancestral sequence. As proteins diverge, destabilizing residues are often mutated to stabilizing residues to offset the effects of mutations that enhance other properties [68]. This tendency means that any destabilizing residues that were present in an ancestral protein are unlikely to be conserved in modern proteins and thus cannot be correctly predicted in a reconstructed ancestor. Stabilizing residues present in an ancestor are likely to be correctly inferred more frequently. This tendency leads to a reconstructed ancestral protein like HNL1 that may be more stable than the actual ancestor.

## Acknowledgements

We thank Fluorescence Innovations, Inc. (Minneapolis, MN) for measuring the melting temperature of HNL1 and Burnell Lauer for measuring heat stability by activity loss.

## Supporting Information

**S1 Fig.**
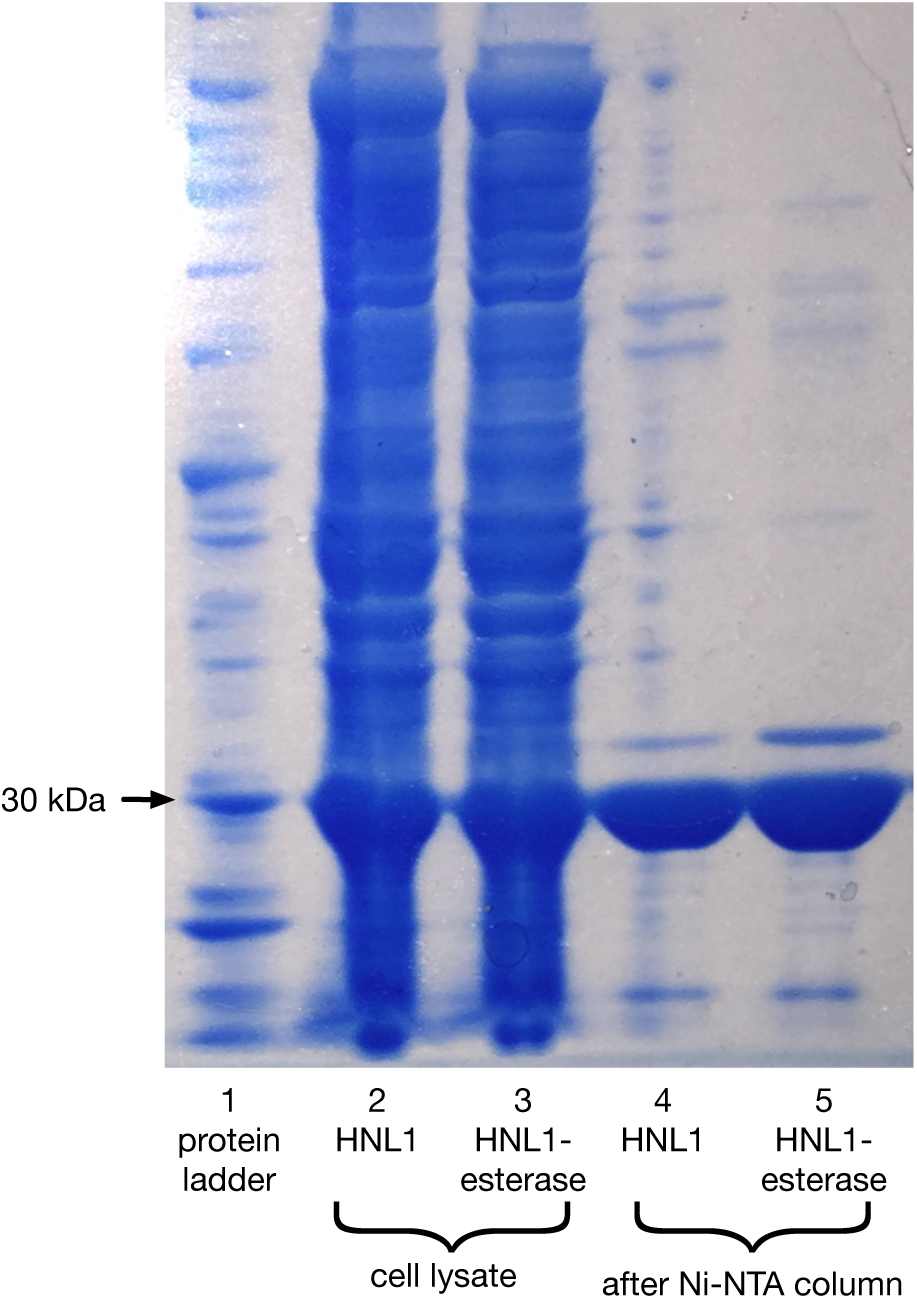
SDS-PAGE gel of HNL1 and HNL1 esterase catalysis variant. Lane 1 is a protein ladder with an arrow marking the 30 kDa band. Lanes 2 and 3 are the cell lysate for HNL1 and HNL1-esterase catalysis variant, respectively. Lanes 4 and 5 are the purified proteins after Ni-NTA chromatography for HNL1 and HNL1-esterase catalysis variant, respectively. The strong bands at ∼30 kDa in lanes 2-5 correspond to the desired proteins (expected molecular weight 31 kDa).

**S2 Fig.**
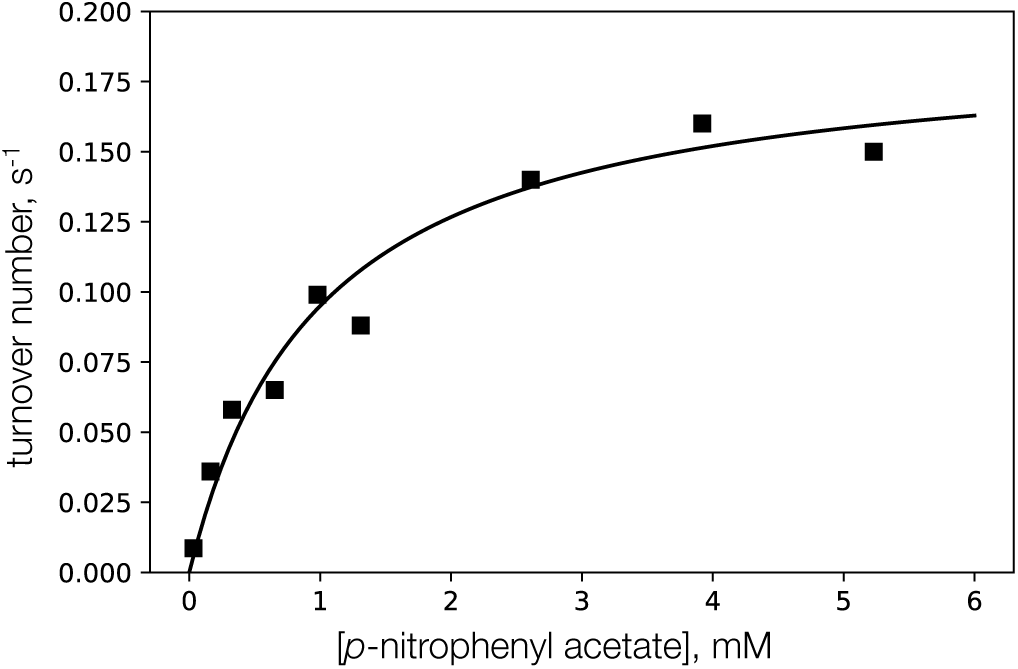
Steady state kinetics for hydrolysis of *pNPAc* catalyzed by the HNL1 esterase catalysis variant. Squares are experimental data and the line is the best fit to the Michaelis-Menten equation where K_M_ = 1.0±0.2 mM and k_cat_ = 0.19±0.01 s^-1^ or 11±1 min^-1^.

**S3 Fig.**
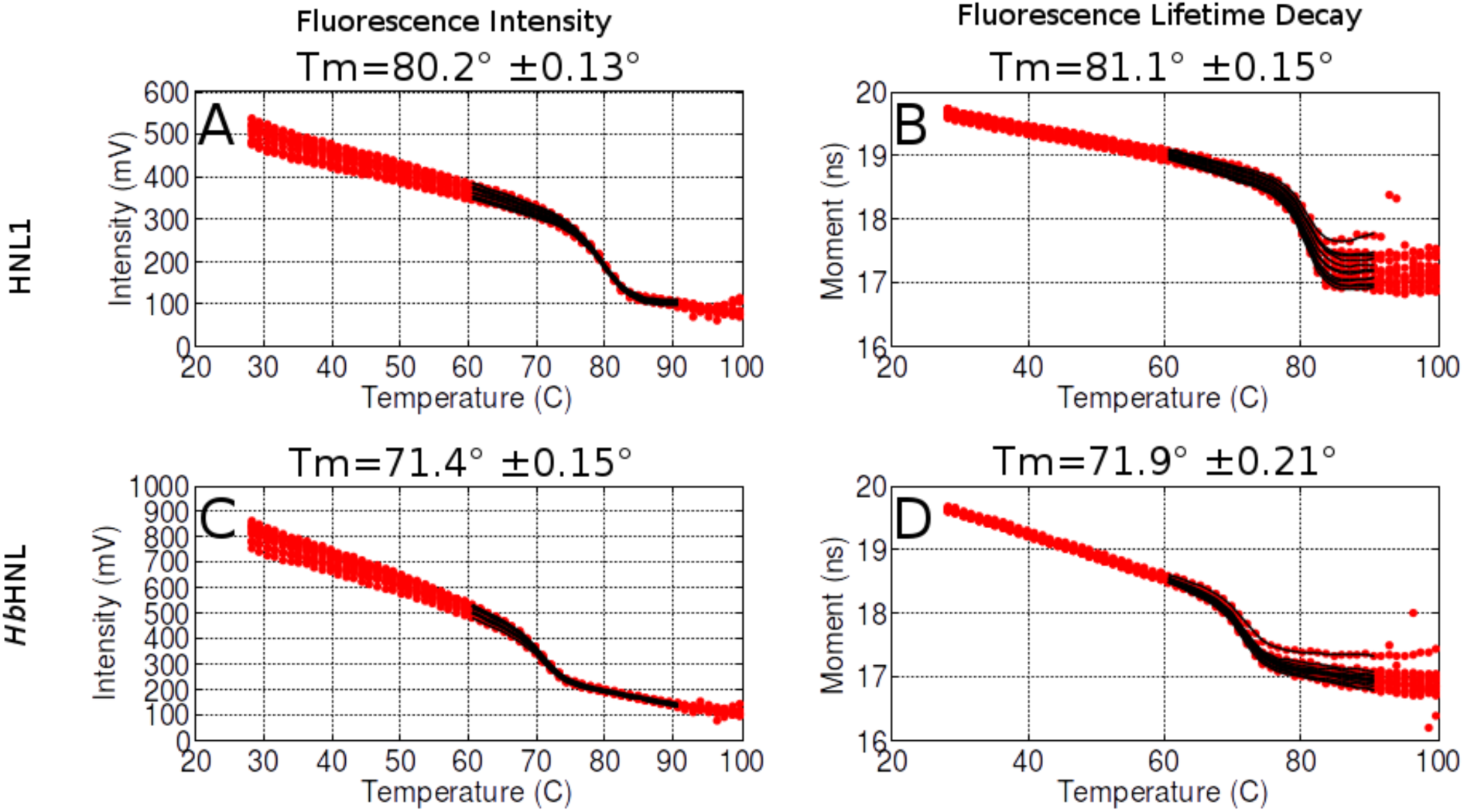
Melting points of HNL1 (A & B) and *Hb*HNL (C & D) measured by decreases in fluorescence intensity (A & C) and decreases in fluorescence lifetime (B & D). The temperature was increased from ambient temperature to 100 °C at 1 °C /min while monitoring the fluorescent properties. The data for 12 replicates of each protein (red) were fit to a theoretical melting curve (black). The average T_m_ and standard deviation are shown above each graph.

**S4 Fig.**
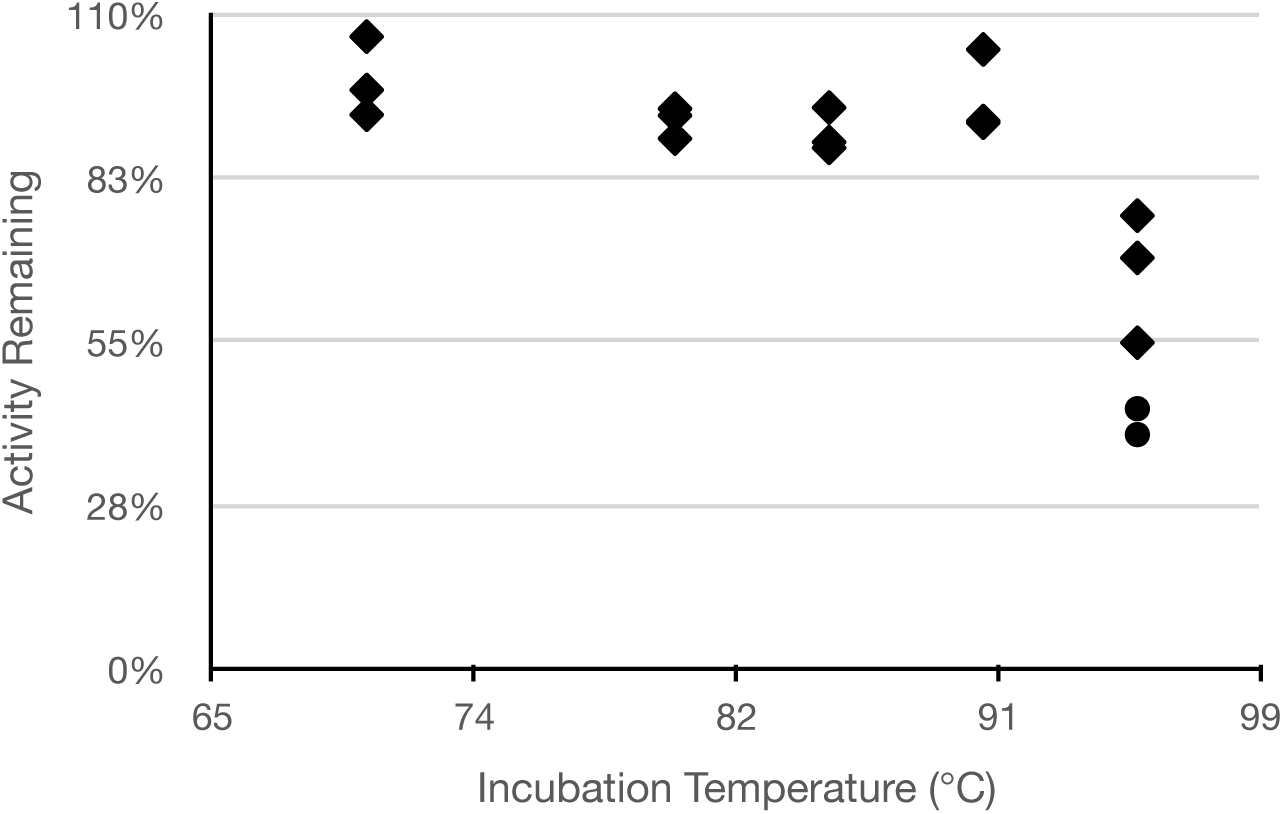
Thermostability of HNL1 to irreversible loss of activity. Samples were heated at the indicated temperature for 15 min (diamonds) or 30 min (circles), cooled on ice for 15 min, then assayed for HNL activity at room temperature. The activity is relative to the activity of unheated samples. Reducing the activity of HNL1 by 50% required 30 min at 95 °C.

**S5 Fig.**
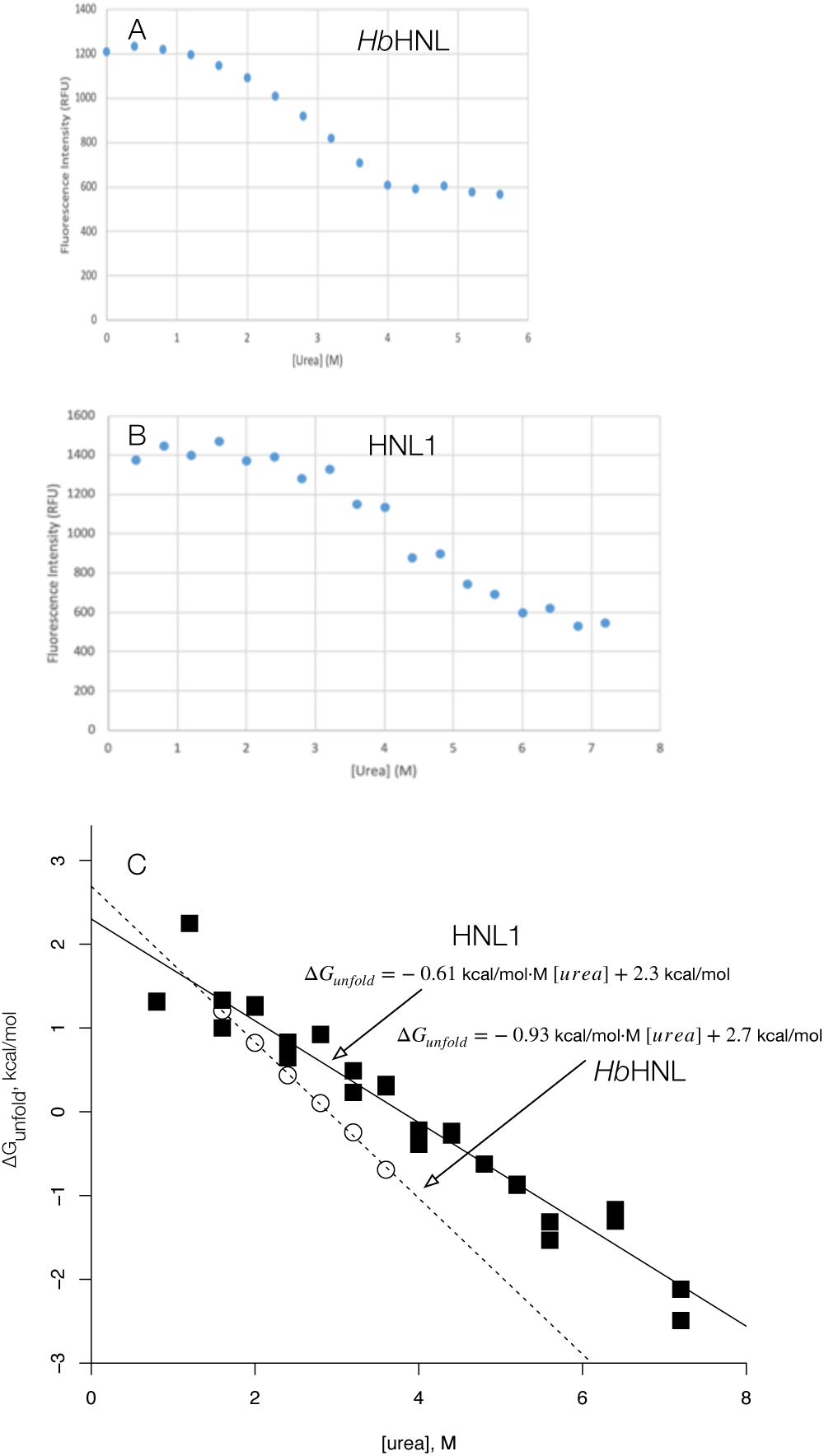
Thermostability of *Hb*HNL and HNL1 measured by urea-induced unfolding. The fluorescence of *Hb*HNL (panel A) and HNL1 (panel B) decreased in solutions of increasing urea concentrations as the proteins unfold. Half of the *Hb*HNL unfolded at 2.8 M urea, while unfolding half of the HNL1 required 3.9 M urea. (C) The slope of the free energy of unfolding versus urea concentration plot is shallower for HNL1 (filled squares, *m*-value = 0.61 kcal/mol·M) than for *Hb*HNL (open circles, m-value = 0.93 kcal/mol·M) indicating that HNL1 unfolds less completely than *Hb*HNL. The y-intercept corresponds to the free energy of unfolding in pure water. Since the slope differs significantly, the difference in the intercepts is not a good measure of difference in stability.

**S6 Fig.**
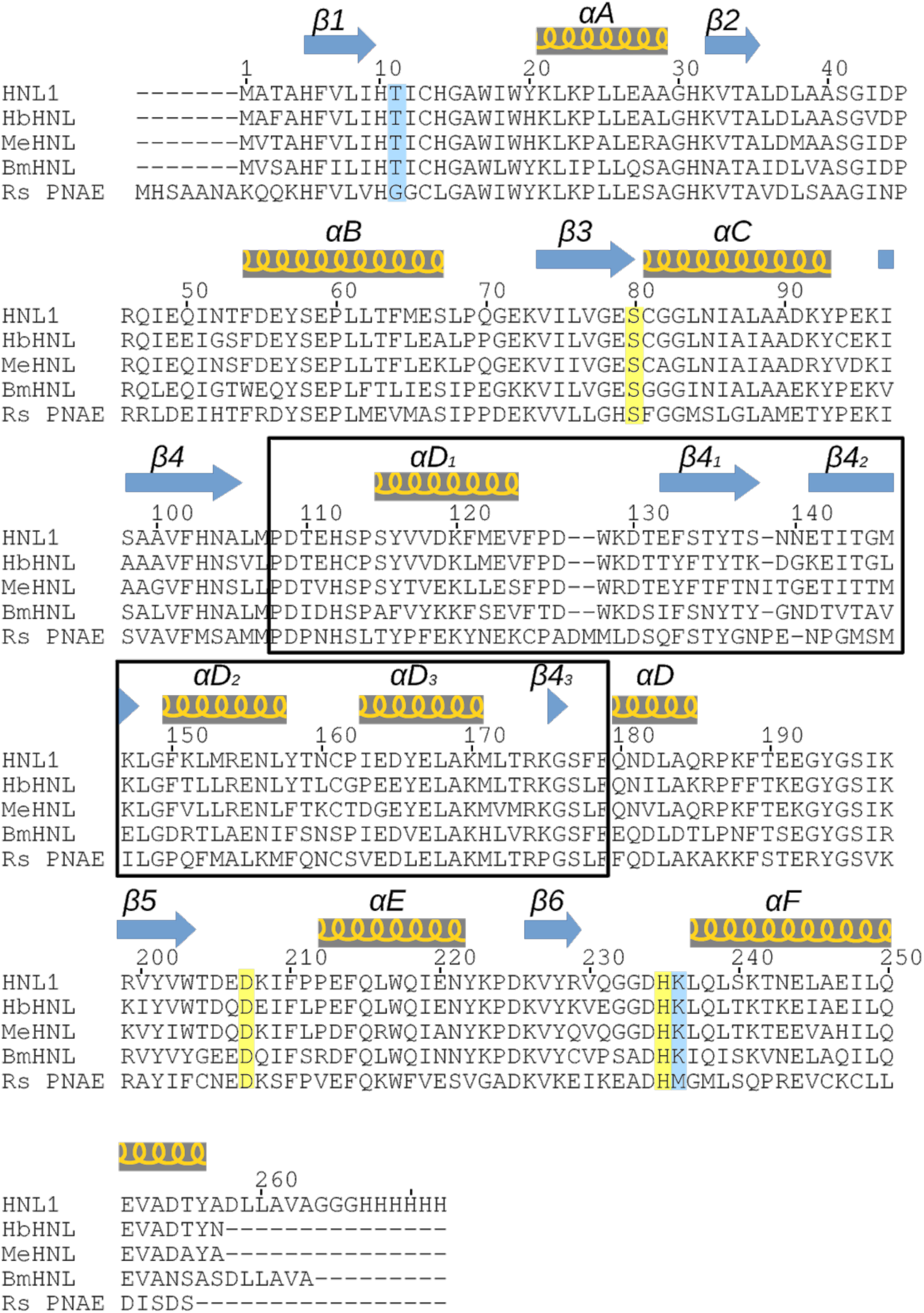
The amino acid sequence of HNL1 aligned to the sequences of the modern proteins *Hb*HNL (P52704.1 pdb id 1yb6), *Me*HNL (P52705.3 pdb id 1dwo), *Bm*HNL (BAI50630.1 pdb id 3wwo), as well as a related esterase from *Rauvolfia serpentina, Rs*PNAE (Q9SE93.1 pdb id 2wfl). Yellow highlights the conserved catalytic triad of Ser, His, and Asp and blue highlights the threonine and lysine, which also contribute to HNL catalysis. Like other esterases, *Rs*PNAE has a glycine in place of threonine and methionine in place of lysine. The cap domain residues are boxed. Secondary structural elements of HNL1 are indicated above alignment and are also similarly positioned in the aligned sequences.

**S7 Fig.**
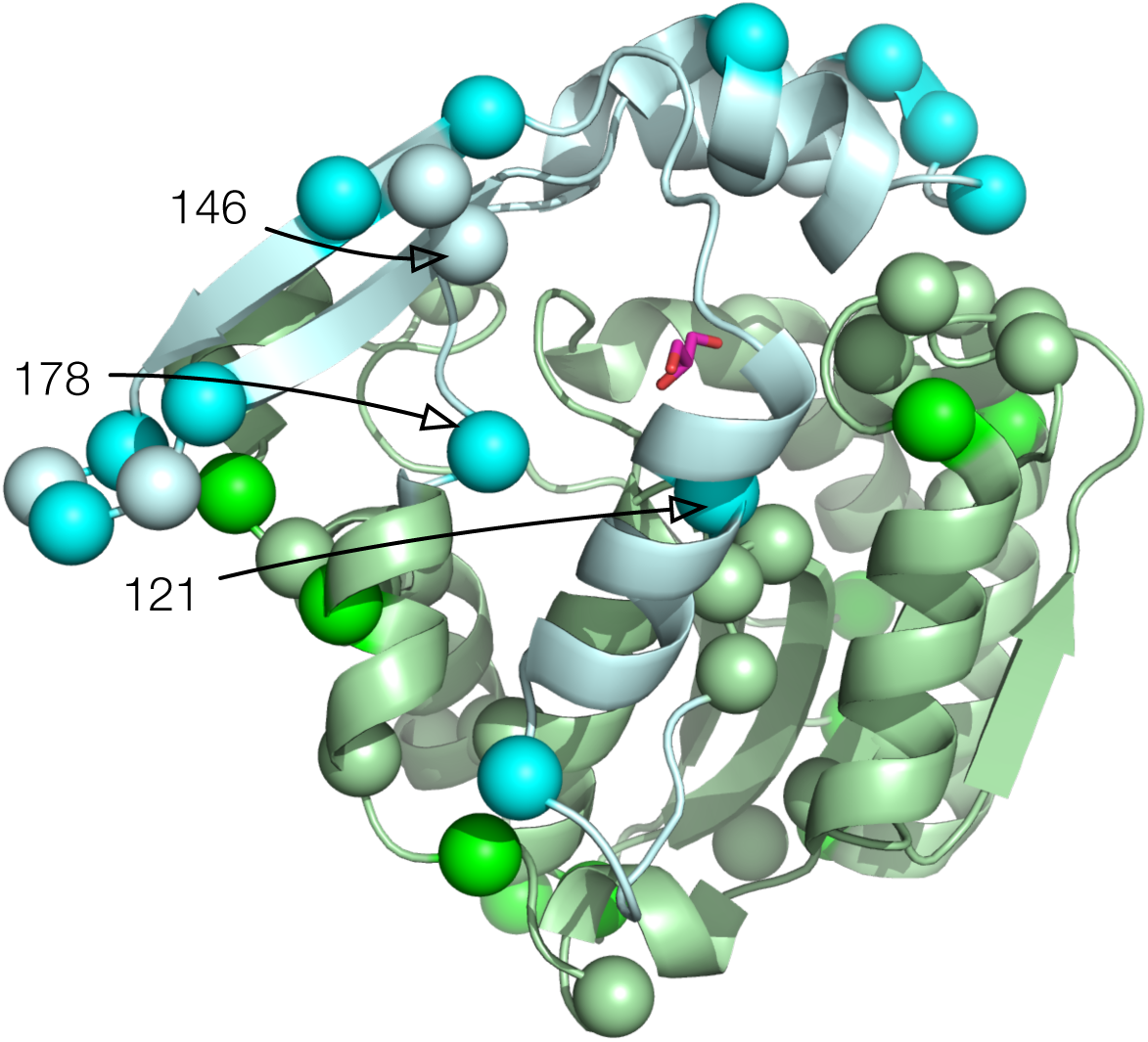
Ribbon diagram of the structure of HNL1 showing the 49 amino acids that differ from HbHNL. Light green marks the catalytic domain, while light cyan marks the lid domain. The Ca atoms of the differing amino acids are shown as spheres. Conservative substitutions (21 in the catalytic domain, 6 in the lid domain) are in the same color as the corresponding ribbon, while non-conservative substitutions (10 in the catalytic domain, 12 in the lid) are in darker colors. The solvent glycerol bound in the active site is in pink and red sticks. The conservative substitutions in the catalytic domain are: 20, 43, 50, 53, 65, 67, 88, 98, 105, 106, 107, 186, 191, 199, 200, 206, 208, 233, 235, 240, 245; the non-conservative substitutions in the catalytic domain are: 3, 30, 52, 70, 94, 182, 188, 211, 243, 257; the conservative substitutions in the lid domain are: 133, 139, 141, 146, 153, 165; the non-conservative substitutions in the lid domain are: 113, 121, 132, 134, 138, 140, 142, 151, 160, 162, 163, 178. The three substitutions responsible for the larger active site in HNL1 in the lid domain are labelled by residue numbers. The substitution at 146 is a conservative substitution from Leu in HbHNL to Met in HNL1. The substitutions at 121 and 178 are non-conservative substitutions from Leu in HbHNL to Phe in HNL1.

**S8 Fig.**
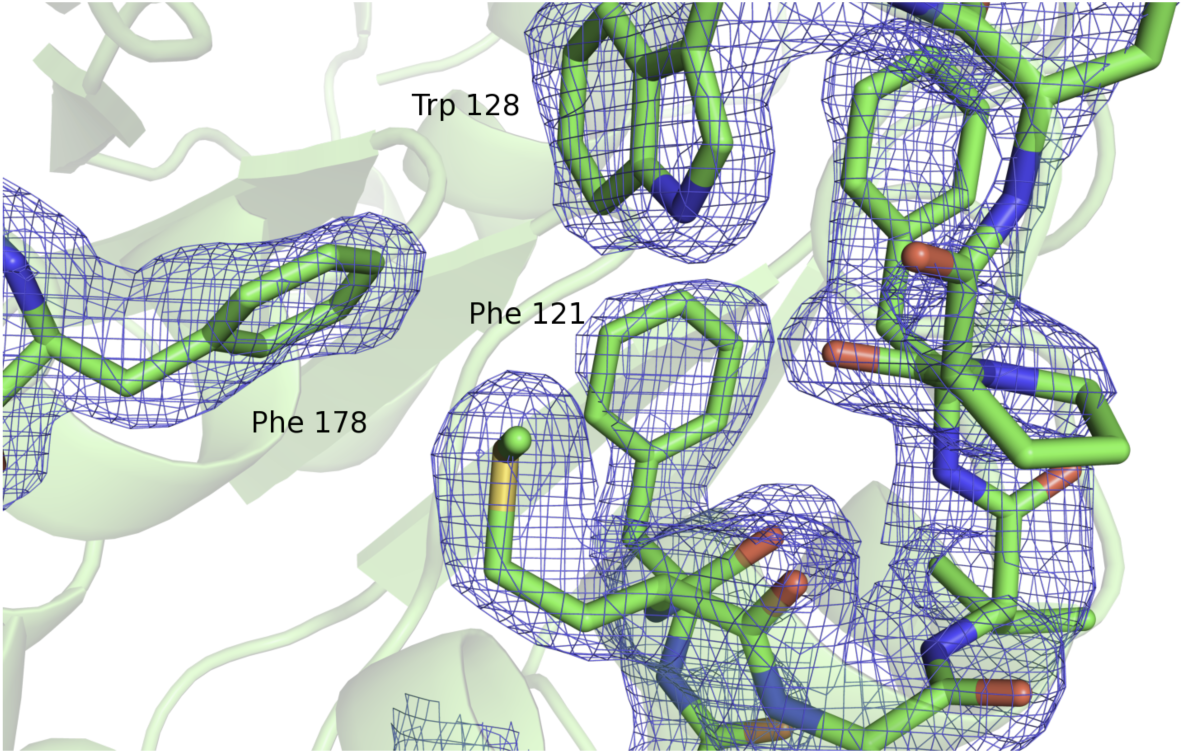
Substrate-binding region of the HNL1 active site showing electron density (2Fo − Fc) contoured at 1 σ. The view is looking into the active site, which is below the plane of the page. Phenylalanines 178 and 121 prevent tryptophan 128 from pushing into the active site. In the modern *Hb*HNL and *Me*HNL, two leucines replace these two phenylalanines.

**S9 Fig.**
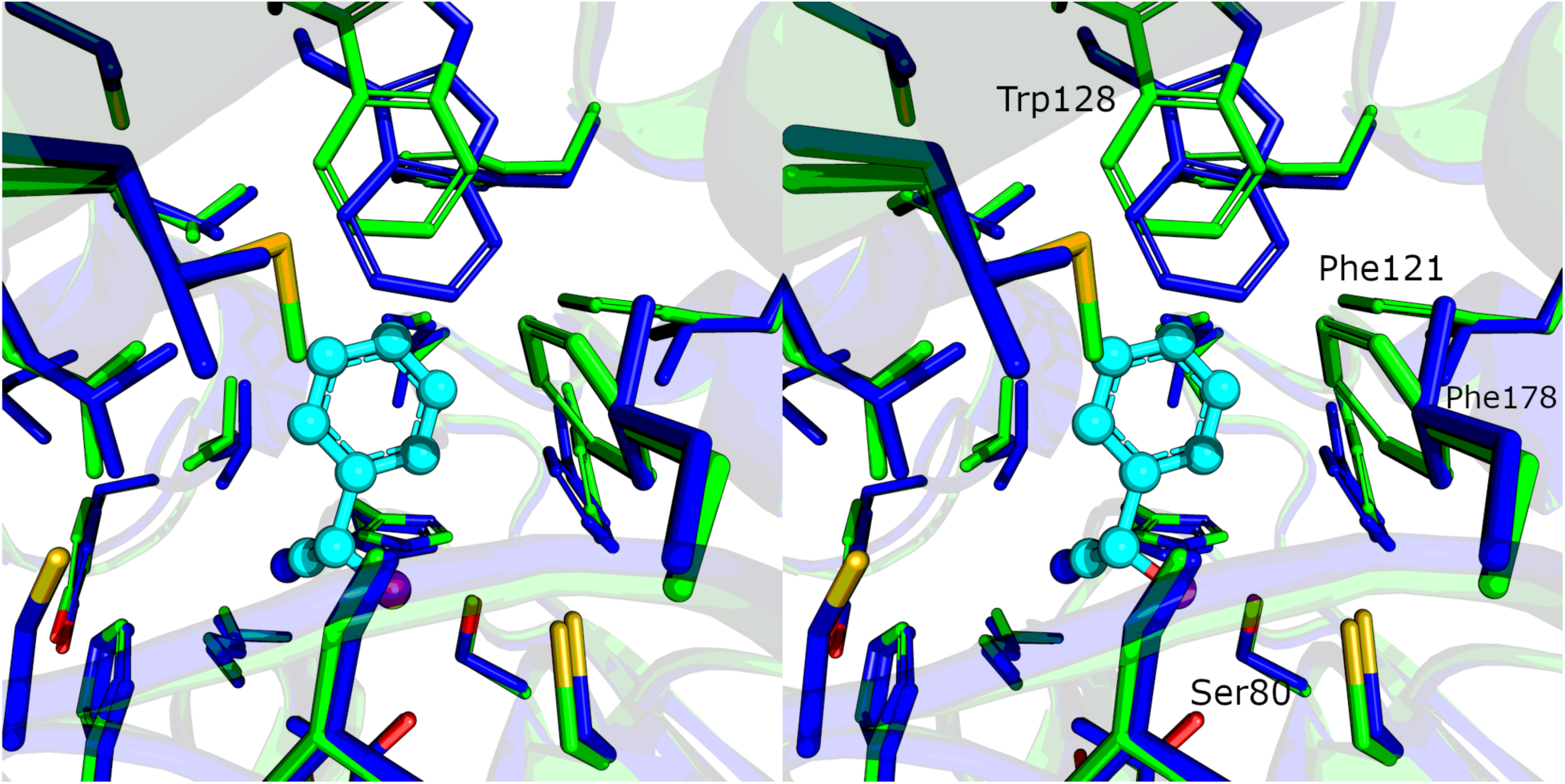
A stereoview of the active sites of HNL1 (pdb id 5tdx, green) and HbHNL (pdb id 3c6x, blue) in sticks representation. Since the structure of HNL1 does not contain bound mandelonitrile, the structure of HbHNL chosen for comparison also lacks bound mandelonitrile. However, mandelonitrile (cyan balls and sticks) has been added as found in another structure of HbHNL (pdb id 1yb6) to orient the viewer.

**S10 Fig.**
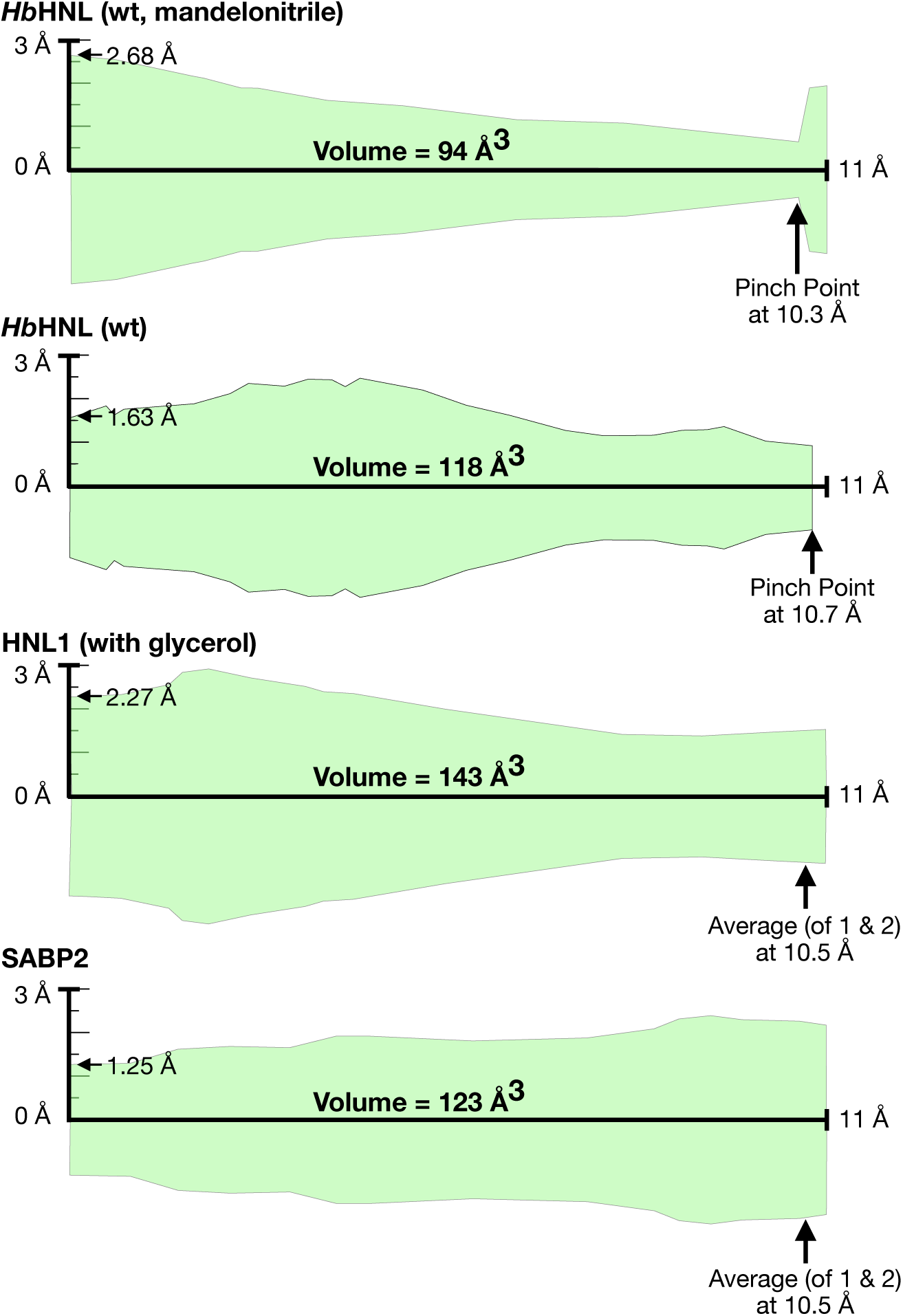
Active-site tunnel cross sections for several enzymes from MOLEonline. The tunnel leading from the active site to the surface for *Hb*HNL with bound mandelonitrile (pdb id 3c6x) shows a pinch point at 10.3 Å, the tunnel for *Hb*HNL without substrate (pdb id 1yb6) shows a pinch point at 10.7 Å, the tunnel for HNL1 shows a narrowing at ∼10.5 Å, and the tunnel for SABP2 with bound product salicylic acid (pdb id 1y7i) shows no narrowing or pinch point. The end of the active site tunnel was defined as 10.5 Å for all enzymes.

**S11 Fig.**
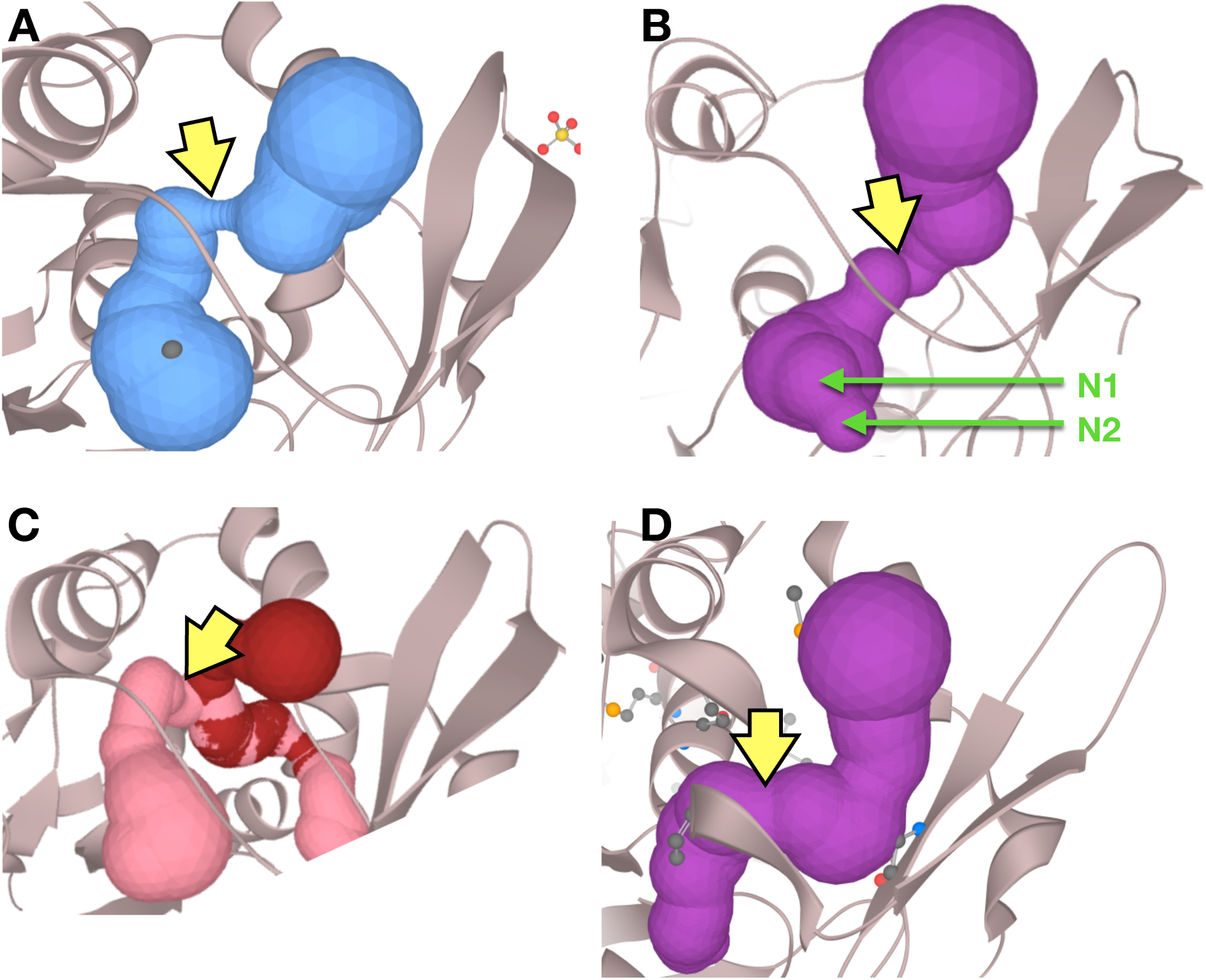
Tunnel containing the active sites for (A) *Hb*HNL with bound mandelonitrile (pdb id 1yb6, MOLEonline tunnel 1), (B) *Hb*HNL without bound substrate (pdb id 3c6x, MOLEonline tunnel 4), (C) HNL1 (pdb id 5tdx, MOLEonline tunnel 11), and (D) SABP2 (pdb id 1y7i, MOLEonline tunnel 4). The same tunnel is compared in each case, although the MOLEonline numbering for each tunnel differed. The start of the active site was defined as the start of the tunnel in *Hb*HNL with bound mandelonitrile. This definition means that for *Hb*HNL without bound substrate, the region labelled N1 in panel B is included, but N2 is not. Yellow arrows indicate the defined end of the active site, based on the consensus pinch points from the bound and unbound *Hb*HNL structures.

**S12 Fig.**
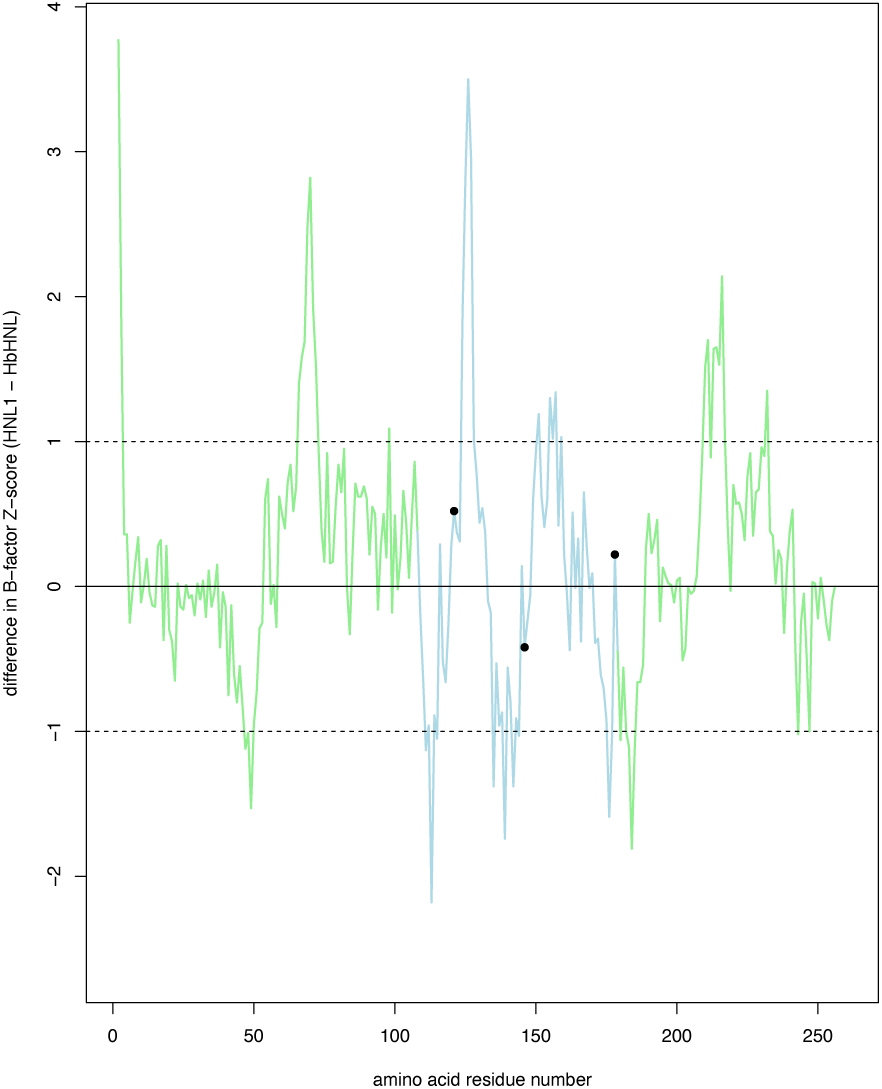
Differences in dynamics of HNL1 (pdb id 5tdx, chain A) and *Hb*HNL (pdb id 1sc9) according to crystallographic B-factors. For each structure the B-factors for the C-α atoms were normalized by conversion to Z-scores to identify regions that are more or less mobile than the average. The Z-scores for the *Hb*HNL were subtracted from those for HNL1 so positive values indicate regions where HNL1 is more mobile than *Hb*HNL and negative values where HNL1 is less mobile. Differences in Z-score greater than 1 or less than -1 (marked by dotted lines) indicate regions where the difference is larger than one standard deviation of the average B-factor. Regions of the HNL1 structure corresponding to these differences are mapped to the structure in Fig 6. The light green lines indicate the catalytic domain, while the light blue line indicates the lid domain. The three substitutions in HNL1 which expand its active site (Phe121, Met146, Phe178) are indicated as black dots on the plot. The Cα atoms that are more mobile in HNL1 by >1 standard deviation are: 2-3, 66-72, 98, 124-127, 151, 155-157, 159, 210-211, 213-216, 232, 254-256. Those less flexible by >1 standard deviation are: 47, 49, 111, 113, 135, 139, 142, 176, 184-185.

**Figure S13.**
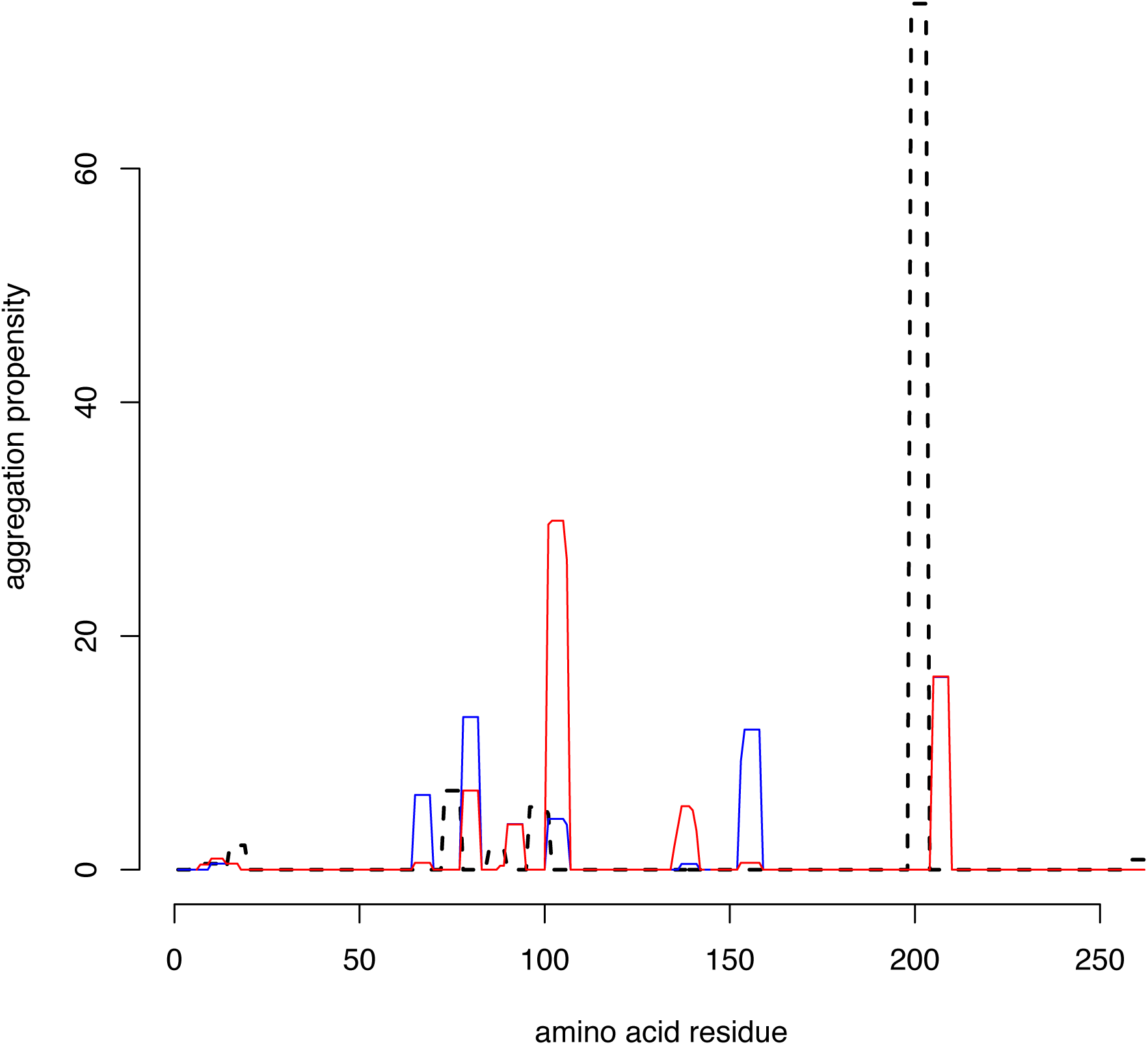
Predicted aggregation propensity of different regions within *Hb*HNL (red line), *Me*HNL (blue line) and HNL1 (dashed black line). All three proteins are predicted to have aggregation-prone regions. Region 200-204 in HNL1 (corresponds to strand β5 in the structure) has the highest predicted aggregation propensity, but experiments showed that HNL1 is less prone to irreversible inactivation than *Hb*HNL and *Me*HNL. Aggregation propensity was calculated using Tango (http://tango.crg.es) using the conditions 72 °C, pH 7, ionic strength 0.02, 1 mM protein.

**S1 Table.**
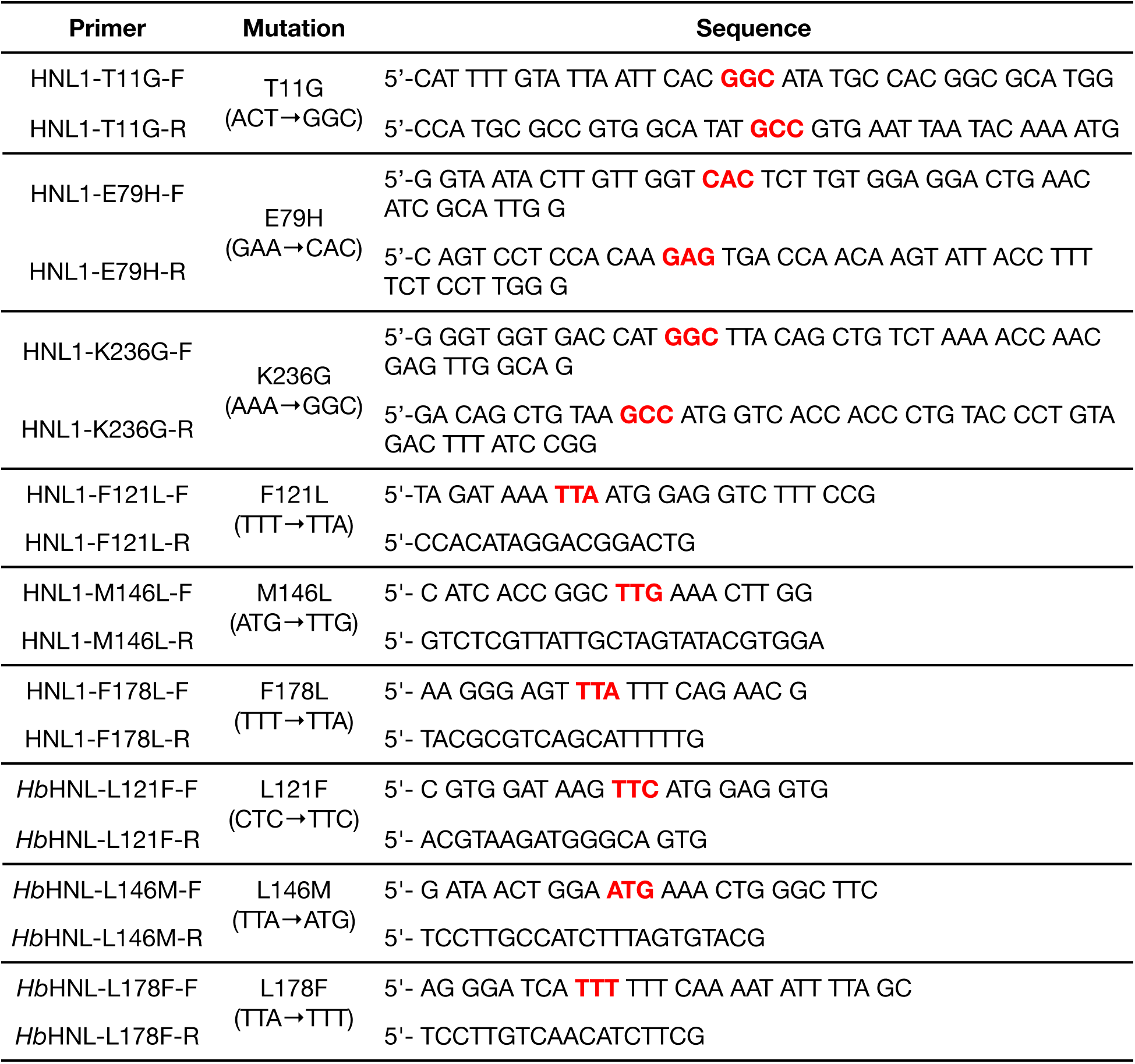
Mutagenic primers for site-directed mutagenesis. The first three primers pairs are overlapping primers for site-directed mutagenesis using the QuickChange (Agilent) method; the last six primer pairs are non-overlapping, back-to-back primers for site-directed mutagenesis using the Q5 (New England Biolabs) method. “F” designates a forward primer and “R” designates a reverse primer. The mutation site is in bold red.

**S2 Table.** Catalytic activity of ancestral enzyme HNL1 and modern enzymes *Hb*HNL and *Me*HNL toward various cyanohydrins and esters and one 2-nitro ethanol.

| Reaction Type | Substrate | HNL1 | HbHNL | MeHNL |
| --- | --- | --- | --- | --- |
| Cyanohydrin cleavage | acetone cyanohydrin | 880 ±70 | 2,400 ±100 | 12,600 ±800<sup>b</sup> |
|  | mandelonitrile | 340 ±10 | 1,530 ±130 | 1,349 ±20 |
|  | lactonitrile<sup>b</sup> | 5.0 ±0.2 | 50 ±11 | 13 ±1 |
|  | 2-hydroxypentanenitrile<sup>b</sup> | 0.5 ±0.1 | 7.2 ±0.8 | 0.66 ±0.06 |
|  | 2-hydroxyhexanenitrile<sup>b</sup> | 0.42 ±0.09 | 24 ±2 | 1.1 ±0.1 |
|  | 2-hydroxy-2-(6-methoxynaphthalen-2-yl) acetonitrile, **1**<sup>c</sup> | 1.06 ±0.04 | 0.64 ±0.05 | Not tested |
| Nitro-aldol cleavage | 2-nitro-1-phenyl ethanol | 14 ±1 | 7.2 ±0.3 | 0.3 |
| Ester hydrolysis | 4-nitrophenyl acetate | 2.5 ±0.1 | 0.19 ±0.06 | 0.060 ±0.005 |
|  | methyl salicylate<sup>d</sup> | 0.048 ±0.006 | 0.066 ±0.006 | <0.09 |
|  | 1-naphthyl acetate<sup>e</sup> | 0.016 ±0.002 | <0.008 | <0.4 |
|  | 2-naphthyl acetate<sup>e</sup> | 0.024 ±0.003 | <0.008 | <0.4 |

<sup>a</sup> Rates are $k_{cat}$ values in min<sup>-1</sup> and taken from Devamani et al. (2016) unless otherwise noted. Errors are standard deviations.
<sup>b</sup> rate at 5 mM substrate.
<sup>c</sup> This work; rate at 0.45 mM substrate.
<sup>d</sup> rate at 0.5 mM substrate.
<sup>e</sup> rate at 2 mM substrate.

**S3 Table.** Location and conservative/non-conservative nature of the 49 amino acid differences between HNL1 and *Hb*HNL.^*a*^

|  | entire protein | lid domain | catalytic domain |
| --- | --- | --- | --- |
| total residues | 257 total<br>188 surface, 73.2% of 257<br>69 interior, 26.8% of 257 | 72 total<br>59 surface, 81.9% of 72<br>13 interior, 18.1% of 72 | 185 total<br>128 surface, 69.2% of 185<br>57 interior, 30.8% of 185 |
| residues that differ | 49, 19.1% of 257<br>40 surface, 81.6% of 49<br>9 interior, 18.4% of 49 | 18, 25% of 72<br>15 surface, 83.3% of 18<br>3 interior, 16.7% of 18 | 31, 16.8% of 185<br>25 surface, 80.6% of 31<br>6 interior, 19.4% of 31 |
| conservative replacements | 27<br>19 surface, 70.4% of 27<br>8 interior, 29.6% of 27 | 6, 33% of 18<br>4 surface, 66.7% of 6<br>2 interior, 33.3% of 6 | 21, 67.7% of 31<br>15 surface, 71.4% of 21<br>6 interior, 28.6% of 21 |
| non-conservative replacements | 22<br>21 surface, 95.5% of 22<br>1 interior, 4.5% of 22 | 12, 66.7% of 18<br>11 surface, 91.7% of 12<br>1 interior, 8.3% of 12 | 10, 32.3% of 31<br>10 surface, 100% of 10<br>0 interior, 0% of 10 |

<sup>a</sup>Surface residues are those that have at least 2.5 Å<sup>2</sup> exposed to the solvent in the HNL1 struc-
ture calculated using the PyMOL script findSurfaceResidues (http://pymolwiki.org/index.php/FindSurfaceResidues).

**S4 Table.** Potential structural contributions to thermostability in *Hb*HNL and HNL1.^*a*^

| property | *HbHNL* | HNL 1 |
| --- | --- | --- |
| Total solvent accessible surface area,<sup>b</sup> Å<sup>2</sup> | 26,722 | 27,024 |
| Non-polar solvent accessible surface area,<sup>c</sup> Å<sup>2</sup> | 16,894 | 16,650 |
| Number of electrostatic interactions<sup>d</sup> | 111 | 109 |
| Number of nonpolar contacts<sup>e</sup> | 256 | 262 |
| Number of hydrogen bonds<sup>f</sup> | 143 | 140 |

<sup>a</sup>Calculations are for one monomer of the protein only without water or bound ligands. *HbHNL* structure pdb id 1yb6 used in comparison.
<sup>b</sup>Calculated using the get\_area function of PyMOL
<sup>c</sup>Non-polar surface area calculated by the exposed surface area of carbon atoms regardless of which residue they occur in.
<sup>d</sup>Number of side chain oxygens in Asp or Glu and side chain nitrogens in Arg, Lys or His with an inter-atomic distance less than 7.0 Å calculated using WHAT IF
<sup>e</sup>Number of interactions between the side chains of Ala, Val, Ile, Leu, Met, Phe, Trp, Pro, or Tyr where the side chain atoms are within 5 Å, Calculated using ProtInter (https://github.com/maxibor/protinter), which is a command line version of PIC: protein interactions calculator.
<sup>f</sup>Calculated optimal hydrogen bond network according to WHAT IF (https://swift.cmbi.umcn.nl/servers/html/index.html).

